# A Computational Approach in Identification of Putative Risk Genes in Parkinsons Disease

**DOI:** 10.1101/2021.10.13.463941

**Authors:** Bhargav Dave, Prasenjit Majumder, Shamayeeta Sarkar

## Abstract

This study focussed on identification of risk genes involved in PD through analysis of microarray data. The two methods were applied viz; WGCNA and DEGs Analysis to identify important genes that are downregulated or upregulated in PD. Both methods show high agreement with each other and also with the available biomedical literature available on this neurodegenerative disease. On the basis of their p–value, 20 significantly upregulated and 19 significantly downregulated genes were found to be playing role in the manifestation of motor and non motor symptoms of the disease, as interpreted from the enrichment analysis.

Gene expression dataset of Parkinson’s Disease (PD) used in this study was obtained from the GeneExpression Omnibus namely GSE8397, GSE20164, and GSE20295 (Edgar, 2002). Among the important genes extracted, the top downregulated and upregulated genes were studied using enrichment analysis. Out of the 19 common downregulated genes, 10 were directly associated with neuron development and differentiation. Two of the genes, FGF13 and CDC42 were associated with multiple signalling pathways. The gene NSF was found to be enriched with GABAergic synapse, associated with the predominating inhibitory neurotransmitter in the mammalian CNS. The inhibitory synapses are thought to provide a brake to neural firing.

The upregulated genes DDIT4, HSPB1, NUPR1, GPNMB and CH13L1 were enriched with apoptotic signalling pathway while MT1M, MT1E, MT1F, MT1X were associated with mineral absorption pathways. Genes like CDC42 have already been reported to be potential diagnostic markers of PD in clinic (Chi et al., 2018).

## 1. Introduction

Parkinsons Disease is one of those neurodegenerative diseases, whose diagnosis and treatment are still to be understood. One of the major challeges is the early diagnosis of PD Though the disease is a slowly progressive disease, with the symptoms worsening over a long period of time, but by the time the symptoms appear, the effect at cellular level is immense and irreparable (Obeso et al., 2017). For example,the death of dopamine neurons in the substantia nigra region of the basal ganglia have been identified to play major role in causing PD but by the time PD symtoms like tremor, rigidity and bradykinesia are visible, there has already been an 80 % loss in dopamine neurons (Obeso et al., 2017).

The symptoms of the disease can be classified into two- motor and non motor. While the former consists of tremors, bradykinesia and muscle rigidity, the latter consists of dementia, pain and loss of smell, to name a few (Greenland and Barker, 2018).

PD is a progressive disease of the Central Nervous System. At cellular level, it involves the loss of dopamine producing cells (Greenland and Barker, 2018).

Hence,it is highly essential to identify early disease diognostic markers (biomarkers) that can be easily studied in clinic through simple tests like blood tests. By definition, “The term biomarker, or biological marker, refers to a broad range of measures which capture what is happening in a cell or organism at a given moment. Biomarkers are objective medical signs (as opposed to symptoms reported by the patient) used to measure the presence or progress of disease” (Califf, 2018).

An observation of the genes whose expressions change significantly in PD as well as in otherwise symptomatically healthy but aged population can be targetted as genes whose expressions progressively change through ageing in potential PD human beings. A further study of such genes in adult population (which are more prone to neurodegenerative diseases like PD), not only in brain but also in blood microarray samples can help in tracking disease onset and progress and used to predict potential diagnostic markers.

## 2. Material and Methods

The following Fig. 1 shows the schematic representation of the workflow of the experiment design.

**Figure 1:**
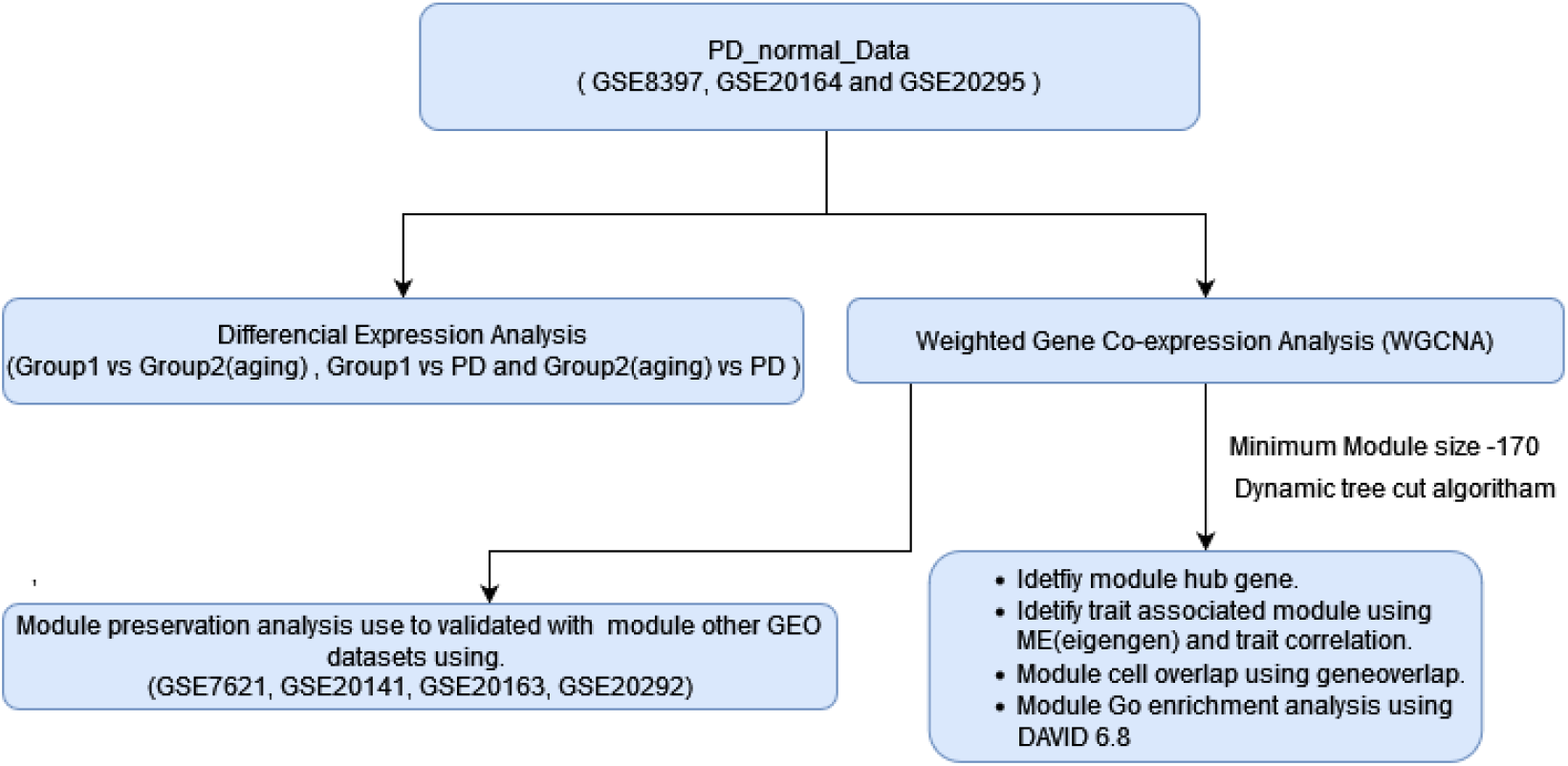
Workflow used to study PD_normal_datasetset

### 2.1. Data Collection and Processing

Microarray dataset of Parkinson’s Disease (PD) was obtained from the Gene Expression Omnibus (GEO) ^1^, a free public functional genomics data repository of an array and sequence-based data (Edgar, 2002). The datasets used in this study were GSE8397, GSE20164, and GSE20295, which are obtained using Affymetrix Human Genome U133A Array chip. The dataset comprised of samples from four brain regions: Substantia Nigra (SN), Prefrontal Cortex (PC), Putamen (PM), Superior Frontal Gyrus (SFG). Samples were between age 41 to 94 years. Depending on the age and sample condition (control/disease), the samples were divided into three groups: Group1 (young), Group2 (aged), and Group3 (PD). Group1 had samples with age between 41 to 70 (less or equal to 70) and neuropathologically healthy individuals, Group2 had samples with age between 70 to 94 (greater than 70) and neuropathologically healthy individuals while Group 3 (PD) consisted of samples with age between 68 to 89 and with Parkinson’s Disease (PD). Additionally four gene expression profiles-GSE7621, GSE20141, GSE20163, GSE20292 were used for preservation analysis. The details of the four datasets which have been used is provided in the Table 1

**Table 1:**
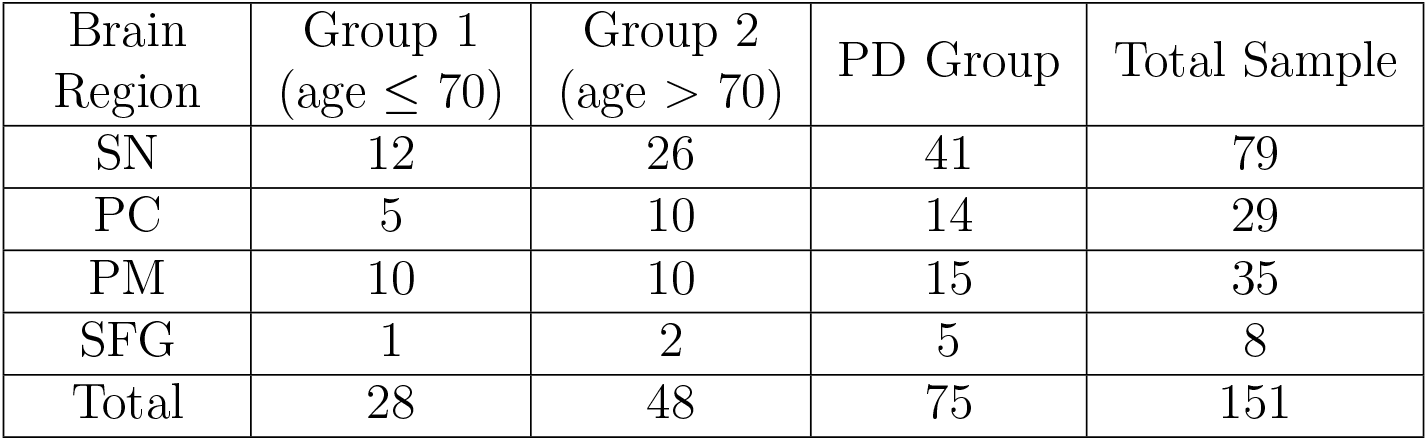
Distribution of samples into group in PD_normal_dataset

The raw data was provided in CEL file for each sample, which were processed. Robust Multichip Average (RMA) procedures was used to process raw data with background correction, normalization, and log transformation.Normalization was done to correct variation between the arrays and within the probe sets. Log transformation was used to improve the distribution of the data. After RMA, The Affy bioconductor package (Gautier et al., 2004) and the Gene Expression Matrix (GEM) or MicroArray Representation “hgu133a.DB” human genome annotation package of R (Carlson) was used to annotate the probe-level data into the gene-level data. Probes with no annotation or with multiple annotation were removed from the analysis. In case of multiple probes associated with the same gene, probes with the highest Interquartile Range (IQR) values was retained. Microarray dataset after preprocessing was represented in the format as shown in Table 2 where the expression values were shown with gene in rows and samples in the column.

**Table 2:**
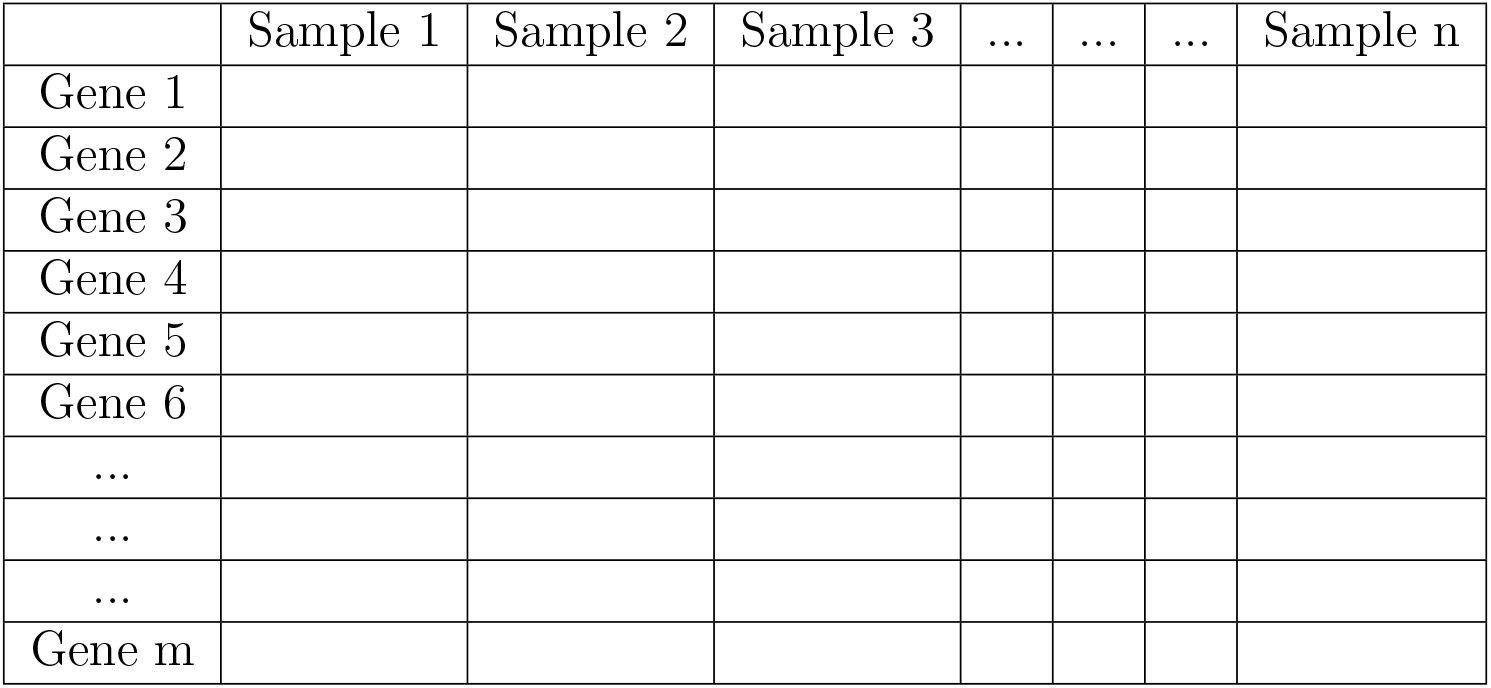
Gene Expression Matrix (GEM) or MicroArray Representation

## 3. Differentially Expressed Gene Analysis

Differentially Expressed Genes, commonly abbreviated as DEGs analysis, is a method of relative determination of changes in expression of one population, relative to another (Lazar et al., 2012) In another way, it mainly focuses on the analysis and interpretation of differences in the gene expression values of the gene between sample group types.

DEGs were determined from the dataset based on the threshold of statistical measure - False discovery rate (FDR) corrected P-value and fold change. FDR corrected P-value was calculated using Benjamini and Hochberg’s method (Ferreira, 2007), and Fold change (FC) difference was calculated (Dalman et al., 2012).The limma package of R (Sakharkar et al., 2019a) was used to find DEGs which provided a facility to calculate FDR corrected P-value and fold change

The genes with positive fold change difference were classified into an upregulated gene, and the genes with negative fold change were classified into a downregulated gene (Lazar et al., 2012). Enrichment analysis of DEGs was done using The Database for Annotation Visualization and Integrated Discovery (DAVID) v6.81. DAVID identifies pathways, biological processes, cellular components, and other functionality of genes(Huang et al., 2008).

## 4. Weighted Gene Co-expression Network Analysis

Weighted Gene Coexpression Network Analysis (WGCNA) is a statistical method for analyzing gene expression data (Zhang and Horvath, 2005). The main focus of WGCNA is to classify genes into a few biologically meaningful modules with similar expression patterns and perform different analyses on the module genes (Horvath, 2011). This was performed using the WGCNA R package (Langfelder and Horvath, 2008). At first, sample-based Hierarchical clustering approach was applied to gene expression data for removing outlier of the sample. Then Pearson correlation was used to generate a gene correlation matrix based random network (Mukaka, 2012).

The genes were classified into modules using a dynamic tree cut algorithm (Langfelder et al., 2007) based on TOM similarity with minimum module size n and then the individual module’s driver gene was determined. Finally, Module enrichment Analysis was done using DAVID version 6.8 (Huang et al., 2008) for each individual module.

The Module cell type overlap was studied to check the significant overlap between the cell-type-specific genes and the module genes. The brain is made up of many cell types, including Astrocytes, Microglia, Oligo-dendrocytes, Neurons, and Endothelial Cells. The overlap between the module and cell-type-specific genes was tested using Fisher’s exact test and a p-value. The GeneOverlap package in R (Shen and Sinai, 2019). (p-value *<* 0.05) was used to study the overlap of a module to the cell type.

### 4.1. Module Preservation Analysis

Preservation of modules is a statistical method used to check the robustness and reproducibility of the defined module across other datasets (Langfelder et al., 2011a). *Z_summary* score and median rank were calculated to measure module preservation.

## 5. Result

### 5.1. Differentially Expressed Gene Analysis

DEGs were obtained from three pair-wise comparison viz. Group1 vs. aging(Group2), Group1 vs. PD, and aging(Group2)vs. PD. Figure 2 shows a Volcano plot, which arranges genes along with biological and statistical significance and helps to decide the threshold of fold change and p-value for a differentially expressed gene. The X-axis gives the log fold change between the two groups, so that up and down-regulation appears symmetrically.Right side of the figure indicate upregulated DEG, and left side indicates down-regulated DEG. The Y-axis represents the FDR corrected p-value comparing samples on a negative log scale, so that expression change with smaller p-values appear higher up. The first one indicates the biological impact of the change, and the second shows the statistical evidence of the change. In this study, a gene with a log fold change greater than 0.5 with a false discovery rate (FDR) corrected p-value less than 0.05 was selected as a differentially expressed gene.

**Figure 2:**
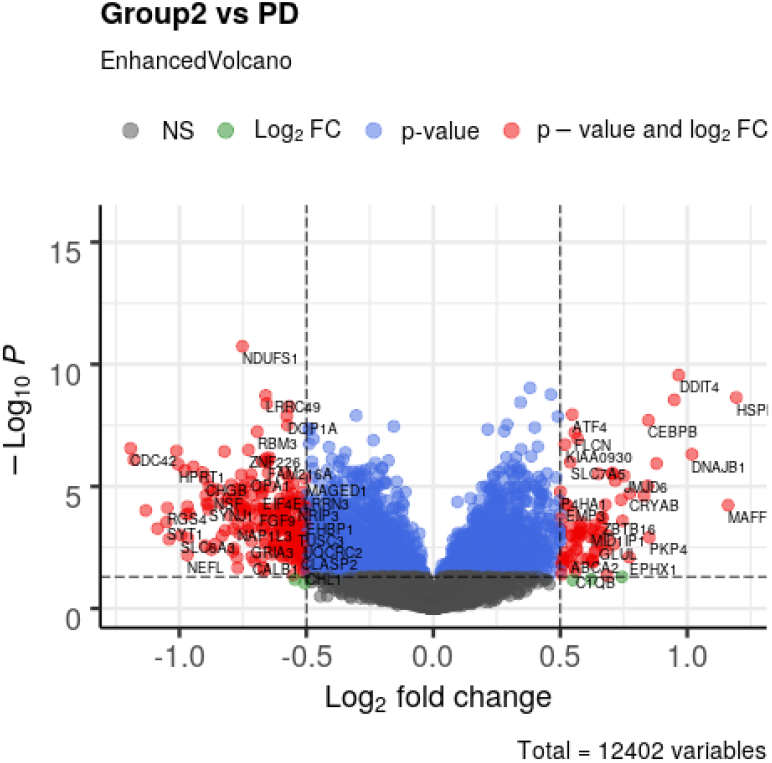
The Volcano plot of Group 2 (aging) vs. PD generated using the EnhancedVolcano R package (Blighe et al., 2019)

From the volcano plot generated using the R package, downregulated and upregulated genes were identified. Table 3, 4, 5, 6 shows the top 25 genes identified under the comparative analysis between Group 1 vs PD and Group 2 vs PD for upregulated and downregulated genes respectively.

**Table 3:**
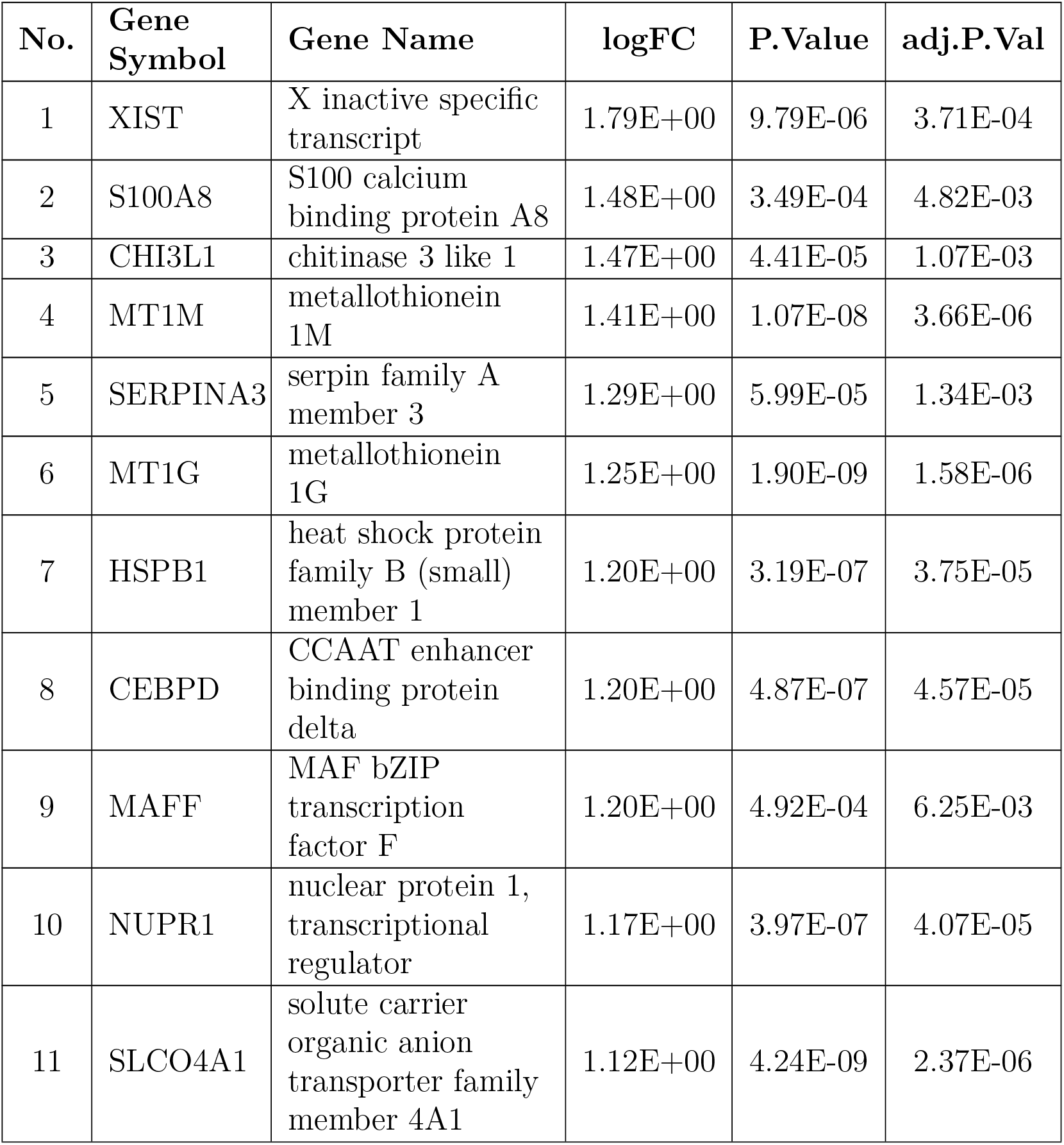
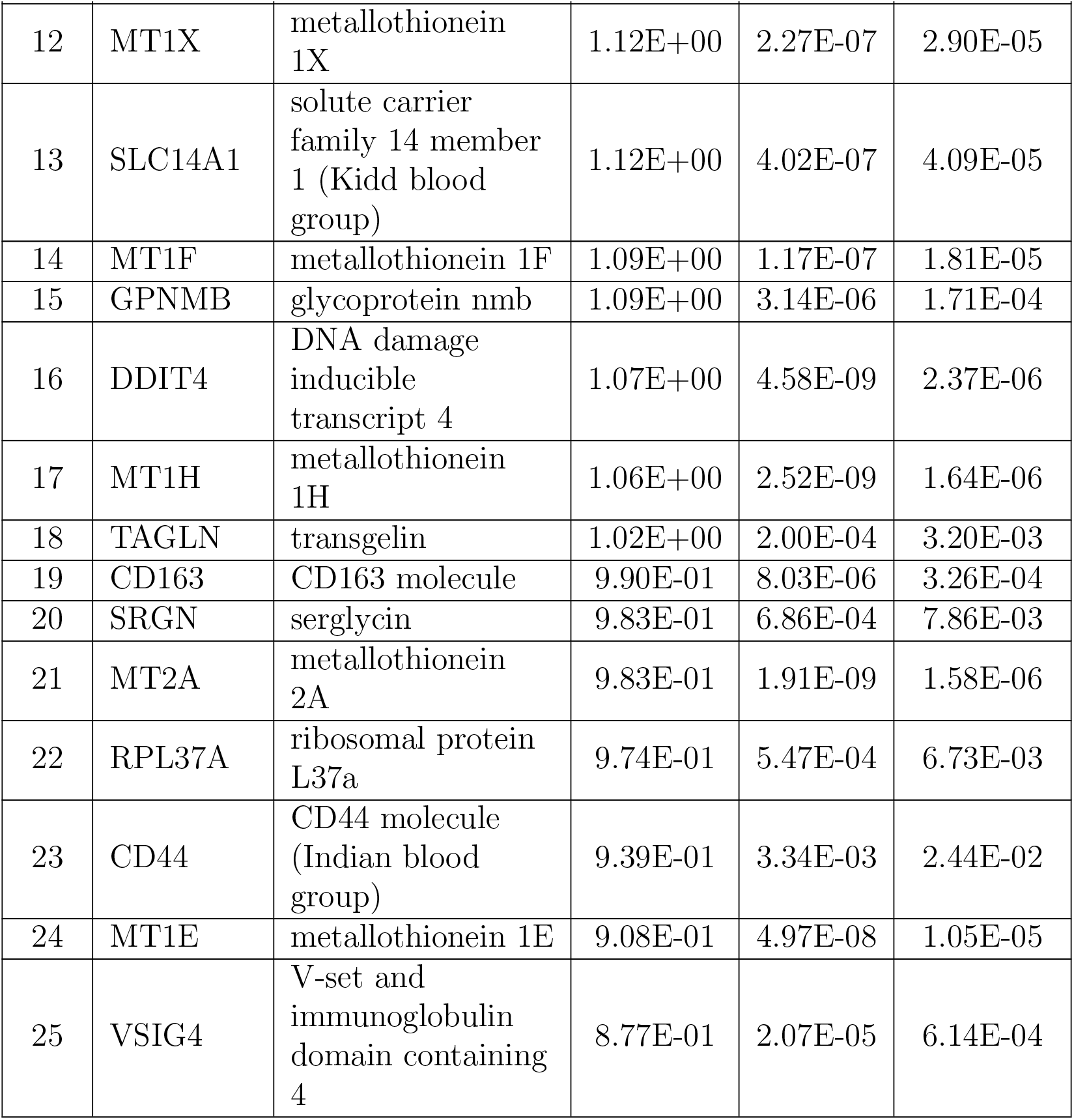
List of top 25 Group1 vs. PD up-regulated genes.

| No. | Gene Symbol | Gene Name | logFC | P.Value | adj.P.Val |
| --- | --- | --- | --- | --- | --- |
| 1 | XIST | X inactive specific transcript | 1.79E+00 | 9.79E-06 | 3.71E-04 |
| 2 | S100A8 | S100 calcium binding protein A8 | 1.48E+00 | 3.49E-04 | 4.82E-03 |
| 3 | CHI3L1 | chitinase 3 like 1 | 1.47E+00 | 4.41E-05 | 1.07E-03 |
| 4 | MT1M | metallothionein 1M | 1.41E+00 | 1.07E-08 | 3.66E-06 |
| 5 | SERPINA3 | serpin family A member 3 | 1.29E+00 | 5.99E-05 | 1.34E-03 |
| 6 | MT1G | metallothionein 1G | 1.25E+00 | 1.90E-09 | 1.58E-06 |
| 7 | HSPB1 | heat shock protein family B (small) member 1 | 1.20E+00 | 3.19E-07 | 3.75E-05 |
| 8 | CEBPD | CCAAT enhancer binding protein delta | 1.20E+00 | 4.87E-07 | 4.57E-05 |
| 9 | MAFF | MAF bZIP transcription factor F | 1.20E+00 | 4.92E-04 | 6.25E-03 |
| 10 | NUPR1 | nuclear protein 1, transcriptional regulator | 1.17E+00 | 3.97E-07 | 4.07E-05 |
| 11 | SLCO4A1 | solute carrier organic anion transporter family member 4A1 | 1.12E+00 | 4.24E-09 | 2.37E-06 |
| 12 | MT1X | metallothionein 1X | 1.12E+00 | 2.27E-07 | 2.90E-05 |
| 13 | SLC14A1 | solute carrier family 14 member 1 (Kidd blood group) | 1.12E+00 | 4.02E-07 | 4.09E-05 |
| 14 | MT1F | metallothionein 1F | 1.09E+00 | 1.17E-07 | 1.81E-05 |
| 15 | GPNMB | glycoprotein nmb | 1.09E+00 | 3.14E-06 | 1.71E-04 |
| 16 | DDIT4 | DNA damage inducible transcript 4 | 1.07E+00 | 4.58E-09 | 2.37E-06 |
| 17 | MT1H | metallothionein 1H | 1.06E+00 | 2.52E-09 | 1.64E-06 |
| 18 | TAGLN | transgelin | 1.02E+00 | 2.00E-04 | 3.20E-03 |
| 19 | CD163 | CD163 molecule | 9.90E-01 | 8.03E-06 | 3.26E-04 |
| 20 | SRGN | serglycin | 9.83E-01 | 6.86E-04 | 7.86E-03 |
| 21 | MT2A | metallothionein 2A | 9.83E-01 | 1.91E-09 | 1.58E-06 |
| 22 | RPL37A | ribosomal protein L37a | 9.74E-01 | 5.47E-04 | 6.73E-03 |
| 23 | CD44 | CD44 molecule (Indian blood group) | 9.39E-01 | 3.34E-03 | 2.44E-02 |
| 24 | MT1E | metallothionein 1E | 9.08E-01 | 4.97E-08 | 1.05E-05 |
| 25 | VSIG4 | V-set and immunoglobulin domain containing 4 | 8.77E-01 | 2.07E-05 | 6.14E-04 |

**Table 4:**
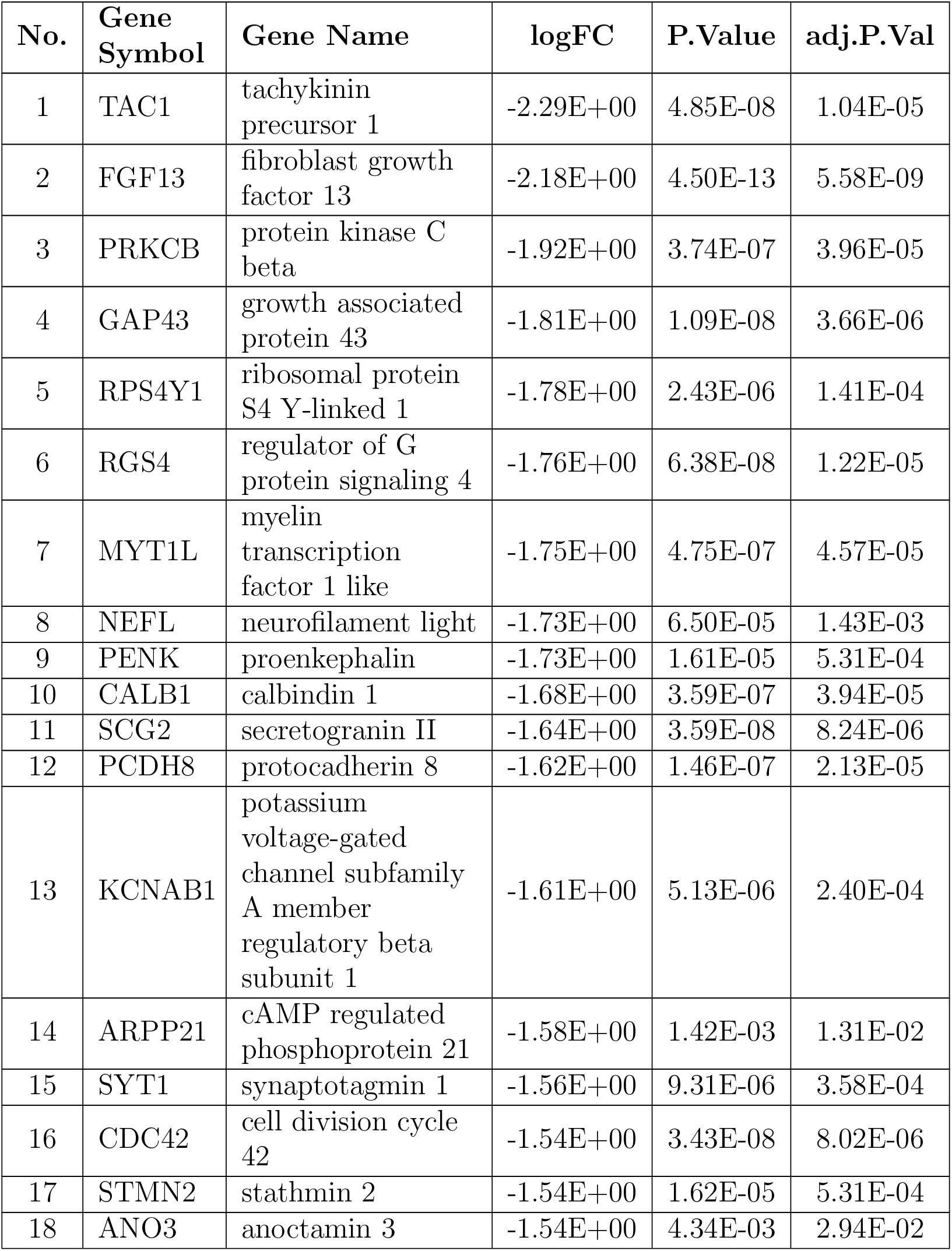
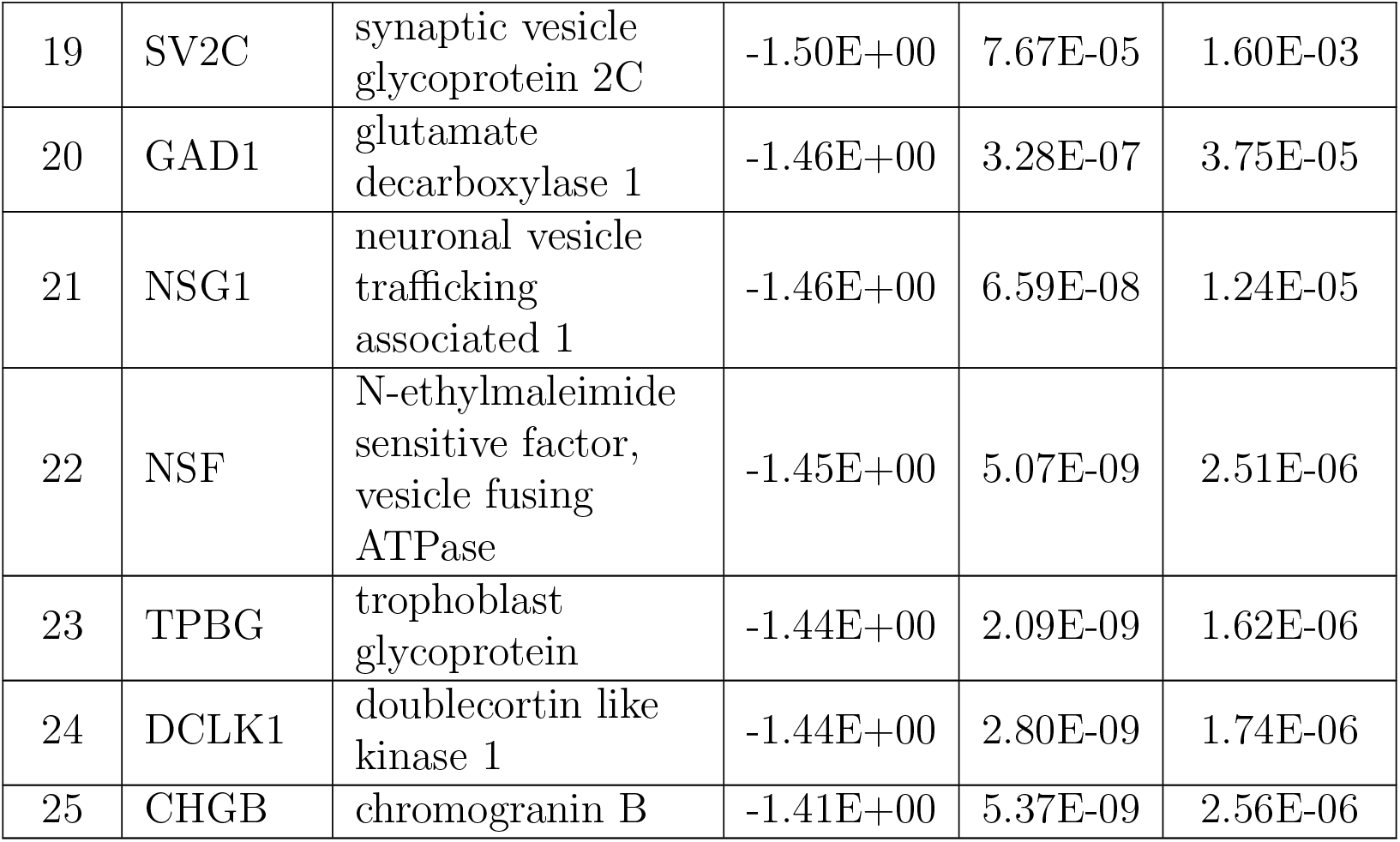
List of top 25 Group1 vs. PD down-regulated genes.

| No. | Gene Symbol | Gene Name | logFC | P.Value | adj.P.Val |
| --- | --- | --- | --- | --- | --- |
| 1 | TAC1 | tachykinin precursor 1 | -2.29E+00 | 4.85E-08 | 1.04E-05 |
| 2 | FGF13 | fibroblast growth factor 13 | -2.18E+00 | 4.50E-13 | 5.58E-09 |
| 3 | PRKCB | protein kinase C beta | -1.92E+00 | 3.74E-07 | 3.96E-05 |
| 4 | GAP43 | growth associated protein 43 | -1.81E+00 | 1.09E-08 | 3.66E-06 |
| 5 | RPS4Y1 | ribosomal protein S4 Y-linked 1 | -1.78E+00 | 2.43E-06 | 1.41E-04 |
| 6 | RGS4 | regulator of G protein signaling 4 | -1.76E+00 | 6.38E-08 | 1.22E-05 |
| 7 | MYT1L | myelin transcription factor 1 like | -1.75E+00 | 4.75E-07 | 4.57E-05 |
| 8 | NEFL | neurofilament light | -1.73E+00 | 6.50E-05 | 1.43E-03 |
| 9 | PENK | proenkephalin | -1.73E+00 | 1.61E-05 | 5.31E-04 |
| 10 | CALB1 | calbindin 1 | -1.68E+00 | 3.59E-07 | 3.94E-05 |
| 11 | SCG2 | secretogranin II | -1.64E+00 | 3.59E-08 | 8.24E-06 |
| 12 | PCDH8 | protocadherin 8 | -1.62E+00 | 1.46E-07 | 2.13E-05 |
| 13 | KCNAB1 | potassium voltage-gated channel subfamily A member regulatory beta subunit 1 | -1.61E+00 | 5.13E-06 | 2.40E-04 |
| 14 | ARPP21 | cAMP regulated phosphoprotein 21 | -1.58E+00 | 1.42E-03 | 1.31E-02 |
| 15 | SYT1 | synaptotagmin 1 | -1.56E+00 | 9.31E-06 | 3.58E-04 |
| 16 | CDC42 | cell division cycle 42 | -1.54E+00 | 3.43E-08 | 8.02E-06 |
| 17 | STMN2 | stathmin 2 | -1.54E+00 | 1.62E-05 | 5.31E-04 |
| 18 | ANO3 | anoctamin 3 | -1.54E+00 | 4.34E-03 | 2.94E-02 |
| 19 | SV2C | synaptic vesicle glycoprotein 2C | -1.50E+00 | 7.67E-05 | 1.60E-03 |
| 20 | GAD1 | glutamate decarboxylase 1 | -1.46E+00 | 3.28E-07 | 3.75E-05 |
| 21 | NSG1 | neuronal vesicle trafficking associated 1 | -1.46E+00 | 6.59E-08 | 1.24E-05 |
| 22 | NSF | N-ethylmaleimide sensitive factor, vesicle fusing ATPase | -1.45E+00 | 5.07E-09 | 2.51E-06 |
| 23 | TPBG | trophoblast glycoprotein | -1.44E+00 | 2.09E-09 | 1.62E-06 |
| 24 | DCLK1 | doublecortin like kinase 1 | -1.44E+00 | 2.80E-09 | 1.74E-06 |
| 25 | CHGB | chromogranin B | -1.41E+00 | 5.37E-09 | 2.56E-06 |

**Table 5:**
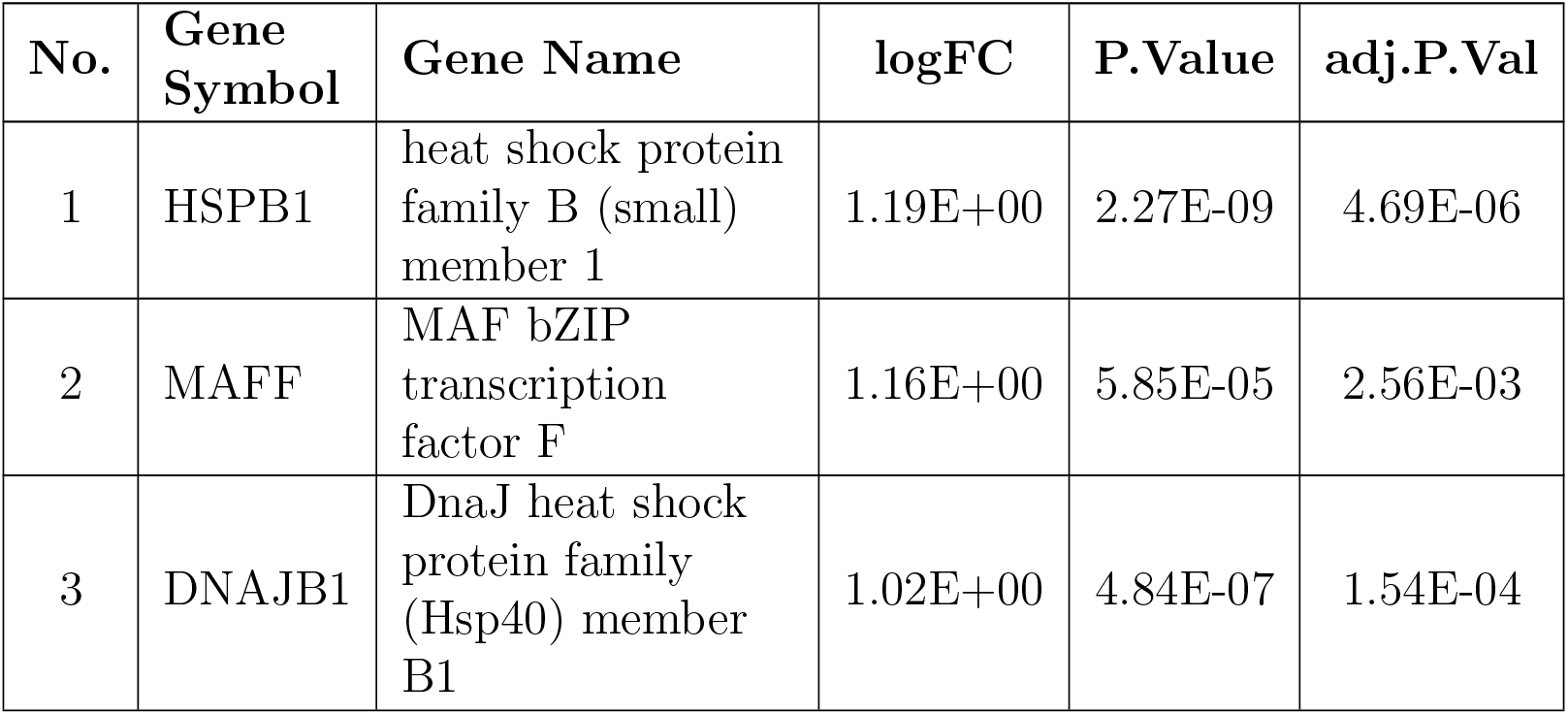
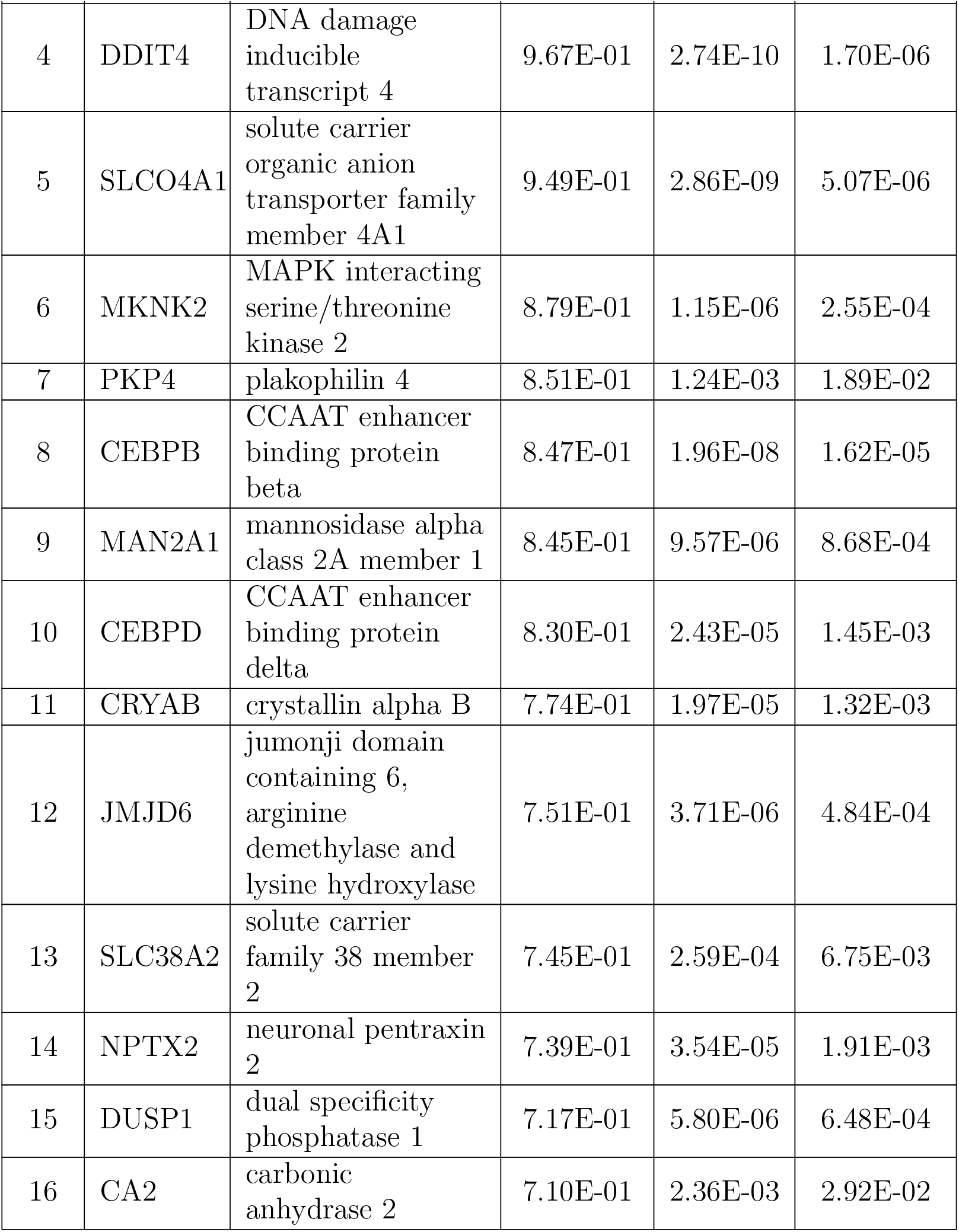
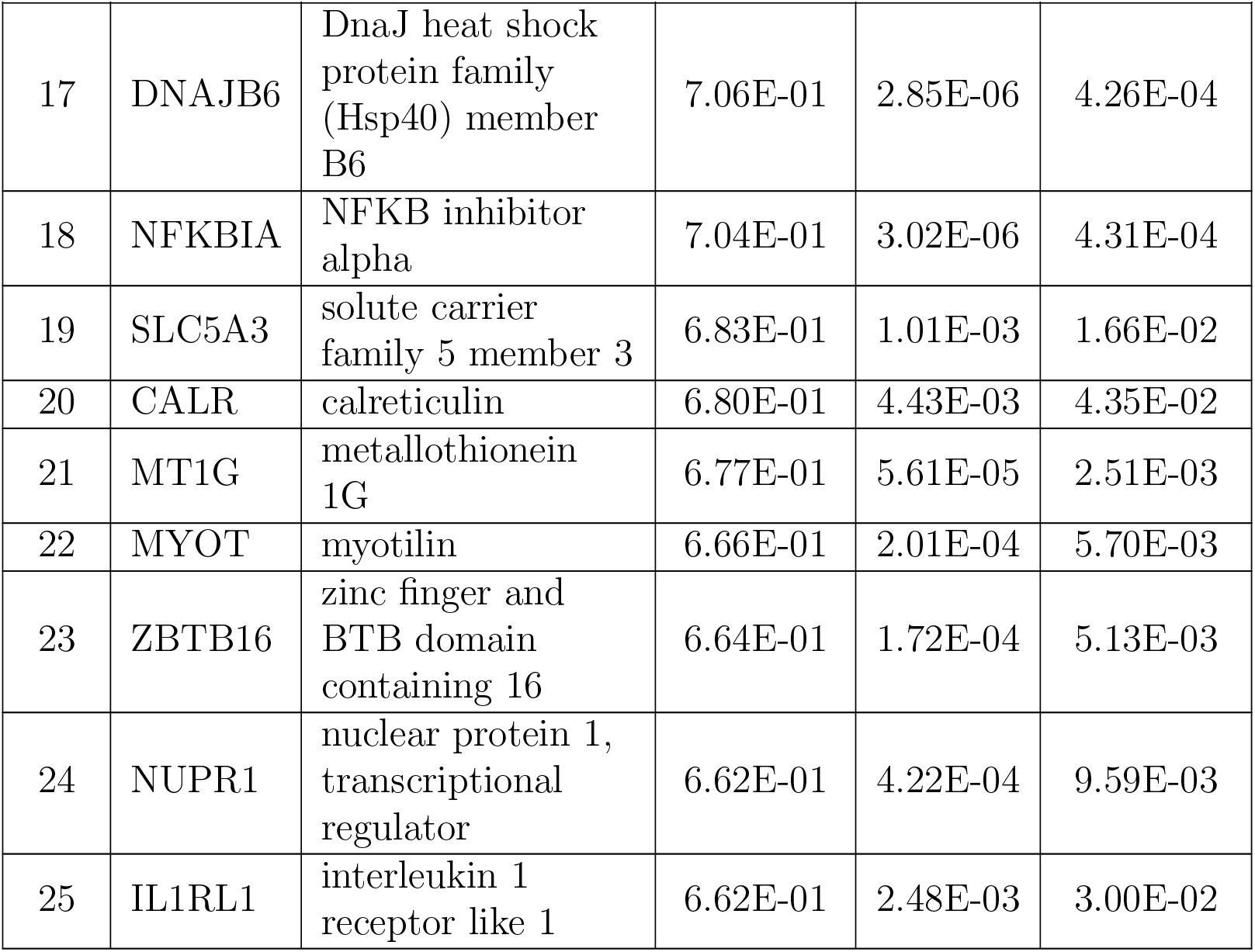
List of top 25 aging (Group2) vs. PD up-regulated genes.

| No. | Gene Symbol | Gene Name | logFC | P.Value | adj.P.Val |
| --- | --- | --- | --- | --- | --- |
| 1 | HSPB1 | heat shock protein family B (small) member 1 | 1.19E+00 | 2.27E-09 | 4.69E-06 |
| 2 | MAFF | MAF bZIP transcription factor F | 1.16E+00 | 5.85E-05 | 2.56E-03 |
| 3 | DNAJB1 | DnaJ heat shock protein family (Hsp40) member B1 | 1.02E+00 | 4.84E-07 | 1.54E-04 |

Continue on the next page
| No. | Gene Symbol | Gene Name | logFC | P.Value | adj.P.Val |
| --- | --- | --- | --- | --- | --- |
| 4 | DDIT4 | DNA damage inducible transcript 4 | 9.67E-01 | 2.74E-10 | 1.70E-06 |
| 5 | SLCO4A1 | solute carrier organic anion transporter family member 4A1 | 9.49E-01 | 2.86E-09 | 5.07E-06 |
| 6 | MKNK2 | MAPK interacting serine/threonine kinase 2 | 8.79E-01 | 1.15E-06 | 2.55E-04 |
| 7 | PKP4 | plakophilin 4 | 8.51E-01 | 1.24E-03 | 1.89E-02 |
| 8 | CEBPB | CCAAT enhancer binding protein beta | 8.47E-01 | 1.96E-08 | 1.62E-05 |
| 9 | MAN2A1 | mannosidase alpha class 2A member 1 | 8.45E-01 | 9.57E-06 | 8.68E-04 |
| 10 | CEBPD | CCAAT enhancer binding protein delta | 8.30E-01 | 2.43E-05 | 1.45E-03 |
| 11 | CRYAB | crystallin alpha B | 7.74E-01 | 1.97E-05 | 1.32E-03 |
| 12 | JMJD6 | jumonji domain containing 6, arginine demethylase and lysine hydroxylase | 7.51E-01 | 3.71E-06 | 4.84E-04 |
| 13 | SLC38A2 | solute carrier family 38 member 2 | 7.45E-01 | 2.59E-04 | 6.75E-03 |
| 14 | NPTX2 | neuronal pentraxin 2 | 7.39E-01 | 3.54E-05 | 1.91E-03 |
| 15 | DUSP1 | dual specificity phosphatase 1 | 7.17E-01 | 5.80E-06 | 6.48E-04 |
| 16 | CA2 | carbonic anhydrase 2 | 7.10E-01 | 2.36E-03 | 2.92E-02 |
| 17 | DNAJB6 | DnaJ heat shock protein family (Hsp40) member B6 | 7.06E-01 | 2.85E-06 | 4.26E-04 |
| 18 | NFKBIA | NFKB inhibitor alpha | 7.04E-01 | 3.02E-06 | 4.31E-04 |
| 19 | SLC5A3 | solute carrier family 5 member 3 | 6.83E-01 | 1.01E-03 | 1.66E-02 |
| 20 | CALR | calreticulin | 6.80E-01 | 4.43E-03 | 4.35E-02 |
| 21 | MT1G | metallothionein 1G | 6.77E-01 | 5.61E-05 | 2.51E-03 |
| 22 | MYOT | myotilin | 6.66E-01 | 2.01E-04 | 5.70E-03 |
| 23 | ZBTB16 | zinc finger and BTB domain containing 16 | 6.64E-01 | 1.72E-04 | 5.13E-03 |
| 24 | NUPR1 | nuclear protein 1, transcriptional regulator | 6.62E-01 | 4.22E-04 | 9.59E-03 |
| 25 | IL1RL1 | interleukin 1 receptor like 1 | 6.62E-01 | 2.48E-03 | 3.00E-02 |

**Table 6:**
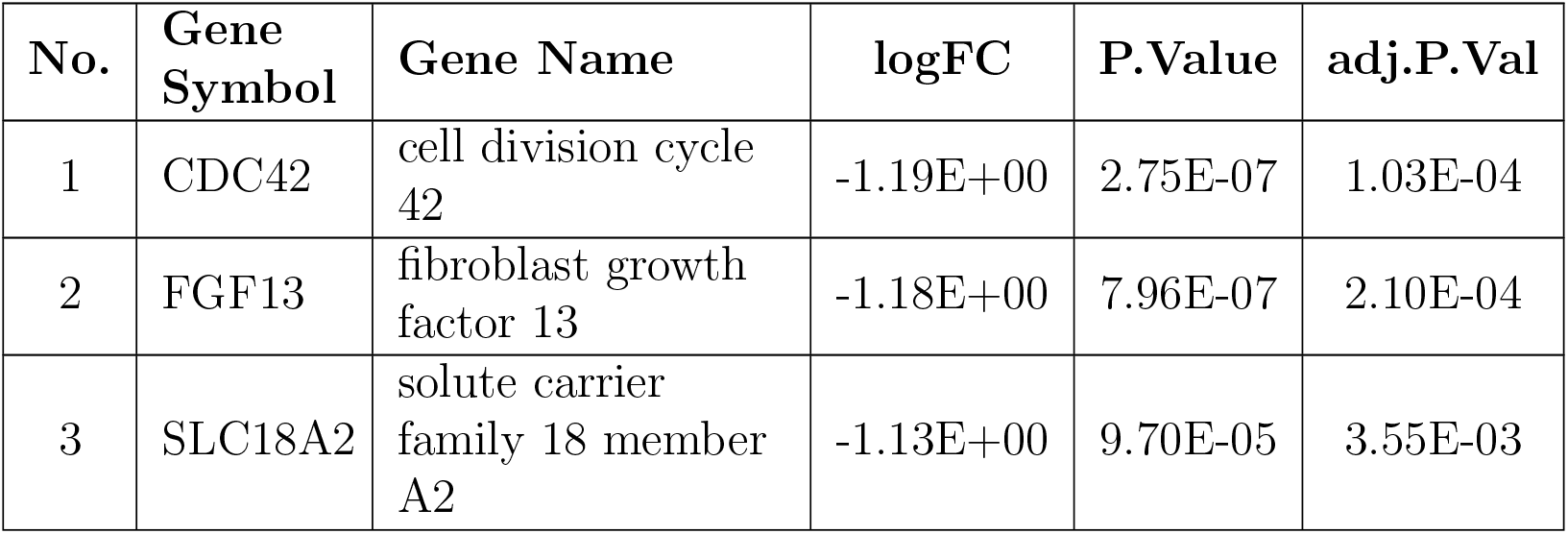
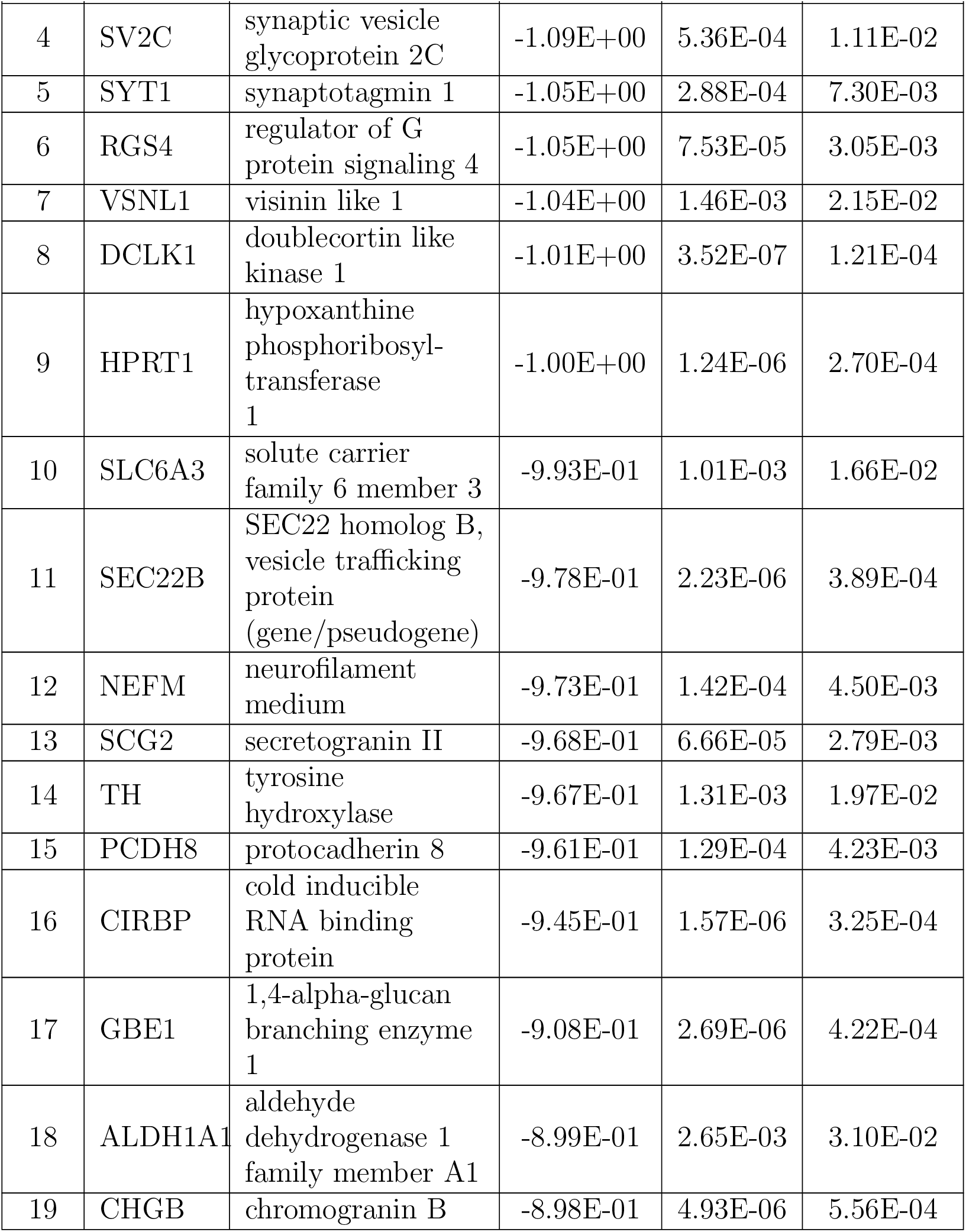
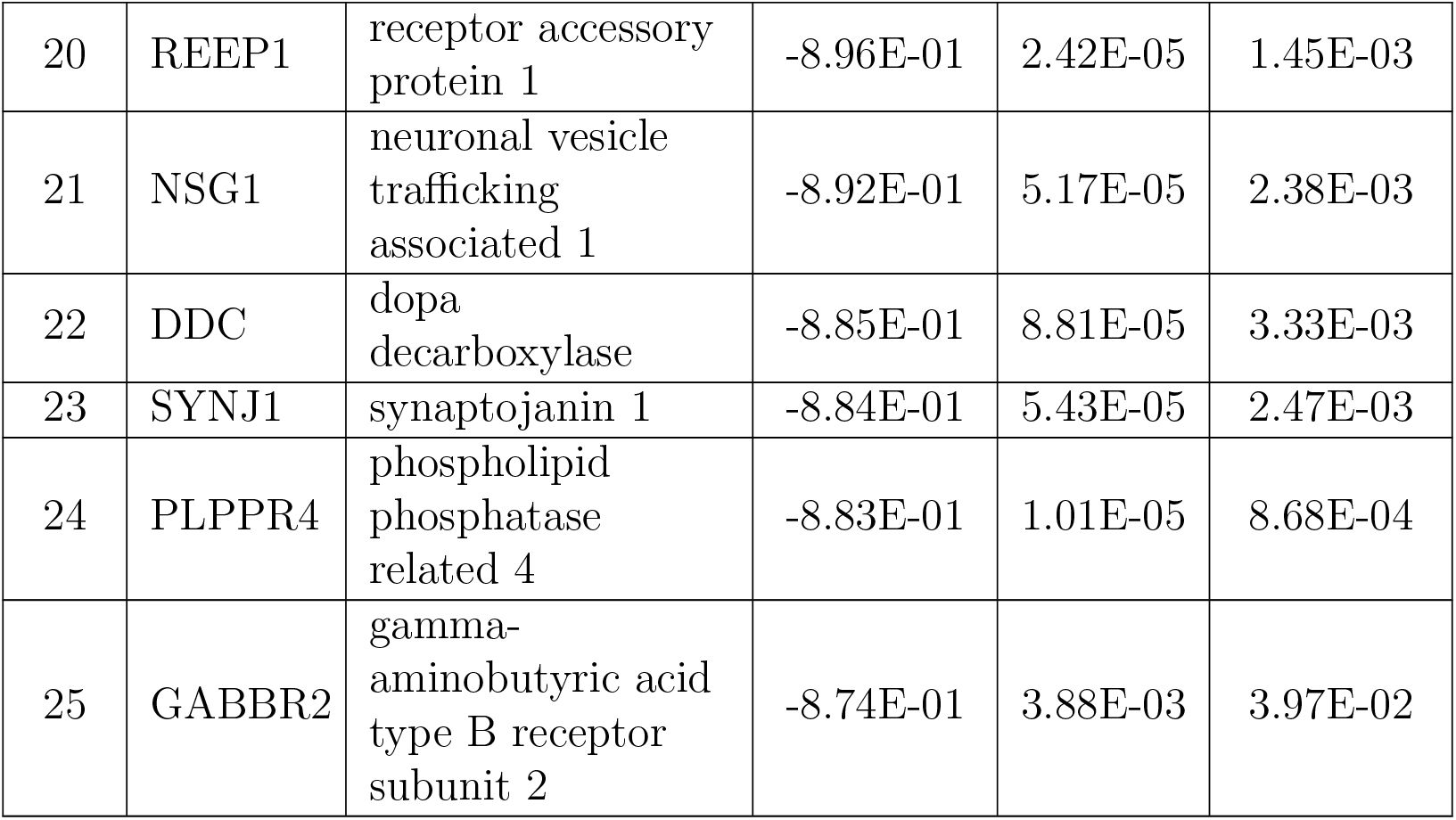
List of top 25 aging (Group2) vs. PD down-regulated genes.

| No. | Gene Symbol | Gene Name | logFC | P.Value | adj.P.Val |
| --- | --- | --- | --- | --- | --- |
| 1 | CDC42 | cell division cycle 42 | -1.19E+00 | 2.75E-07 | 1.03E-04 |
| 2 | FGF13 | fibroblast growth factor 13 | -1.18E+00 | 7.96E-07 | 2.10E-04 |
| 3 | SLC18A2 | solute carrier family 18 member A2 | -1.13E+00 | 9.70E-05 | 3.55E-03 |

Continue on the next page
| No. | Gene Symbol | Gene Name | logFC | P.Value | adj.P.Val |
| --- | --- | --- | --- | --- | --- |
| 4 | SV2C | synaptic vesicle glycoprotein 2C | -1.09E+00 | 5.36E-04 | 1.11E-02 |
| 5 | SYT1 | synaptotagmin 1 | -1.05E+00 | 2.88E-04 | 7.30E-03 |
| 6 | RGS4 | regulator of G protein signaling 4 | -1.05E+00 | 7.53E-05 | 3.05E-03 |
| 7 | VSNL1 | visinin like 1 | -1.04E+00 | 1.46E-03 | 2.15E-02 |
| 8 | DCLK1 | doublecortin like kinase 1 | -1.01E+00 | 3.52E-07 | 1.21E-04 |
| 9 | HPRT1 | hypoxanthine phosphoribosyl-transferase 1 | -1.00E+00 | 1.24E-06 | 2.70E-04 |
| 10 | SLC6A3 | solute carrier family 6 member 3 | -9.93E-01 | 1.01E-03 | 1.66E-02 |
| 11 | SEC22B | SEC22 homolog B, vesicle trafficking protein (gene/pseudogene) | -9.78E-01 | 2.23E-06 | 3.89E-04 |
| 12 | NEFM | neurofilament medium | -9.73E-01 | 1.42E-04 | 4.50E-03 |
| 13 | SCG2 | secretogranin II | -9.68E-01 | 6.66E-05 | 2.79E-03 |
| 14 | TH | tyrosine hydroxylase | -9.67E-01 | 1.31E-03 | 1.97E-02 |
| 15 | PCDH8 | protocadherin 8 | -9.61E-01 | 1.29E-04 | 4.23E-03 |
| 16 | CIRBP | cold inducible RNA binding protein | -9.45E-01 | 1.57E-06 | 3.25E-04 |
| 17 | GBE1 | 1,4-alpha-glucan branching enzyme 1 | -9.08E-01 | 2.69E-06 | 4.22E-04 |
| 18 | ALDH1A1 | aldehyde dehydrogenase 1 family member A1 | -8.99E-01 | 2.65E-03 | 3.10E-02 |
| 19 | CHGB | chromogranin B | -8.98E-01 | 4.93E-06 | 5.56E-04 |
| 20 | REEP1 | receptor accessory protein 1 | -8.96E-01 | 2.42E-05 | 1.45E-03 |
| 21 | NSG1 | neuronal vesicle trafficking associated 1 | -8.92E-01 | 5.17E-05 | 2.38E-03 |
| 22 | DDC | dopa decarboxylase | -8.85E-01 | 8.81E-05 | 3.33E-03 |
| 23 | SYNJ1 | synaptojanin 1 | -8.84E-01 | 5.43E-05 | 2.47E-03 |
| 24 | PLPPR4 | phospholipid phosphatase related 4 | -8.83E-01 | 1.01E-05 | 8.68E-04 |
| 25 | GABBR2 | gamma-aminobutyric acid type B receptor subunit 2 | -8.74E-01 | 3.88E-03 | 3.97E-02 |

### 5.2. WGCNA (Weighted Gene Co-Expression Network Analysis

The PD_normal_datasetset generated upon prepossessing and annotation steps contain 12402 genes and 151 samples. Gene with IQR *>* 0.2 across the 3 groups (Group 1, Group 2 (aged) and PD) were considered for WGCNA analysis to remove non-varying genes as well as decrease computation time (Oldham et al., 2006; Zhang and Horvath, 2005). This screening resulted in 11104 genes. Sample outlier were detected and removed using the sample based hierarchial clustering method (Zhang and Horvath, 2005). Figure (B.9) 3 displays the hierarchical clustering dendrogram, based on which the sample was generated using PD_normal_dataset. Gene similarity matrix was calculated using Pearson correlation (Mukaka, 2012) and transformed to the adjacency matrix using powerlow (Broido and Clauset, 2019). The Figure 4 shows the network achieved scale-free topology at ten. Hence, a soft power (*β*) value of 10 was used to construct a signed co-expression network. The hierarchical clustering, based on the gene with TOM similarity (Yip and Horvath, 2007), generated the tree-type structure Figure 5 with a minimum module size of 170). The Figure 6 shows dendrogram based on the genes generated with modules colors (turquoise, blue, brown, yellow, green, red, black, and pink). A total of 690 outlier genes were detected (grey color). Furthermore, the individual module’s hub (Driver) gene was determined, which showed the highest connectivity in the module. The following driver or hub genes, namely PCMT1, CAPZB, IGF2-AS, NFAT5, PRB4, HPRT1, ATP5MC3, TYROBP, C1orf105, FUT7, RHOT1, RSL24D1, USP33, ATP11B, SSX2IP were found. The Table 7 provides details of all modules with driver genes associated with the individual module.

**Table 7:** Details of the module size and drive or hub gene that generated using PD_normal_dataset.

| Moduel | Moduel Color | Module Size | Driver gene gene |
| --- | --- | --- | --- |
| 0 | gray | 690 | - |
| 1 | turquoise | 1847 | ATP11B |
| 2 | blue | 1407 | CAPZB |
| 3 | brown | 1028 | IGF2-AS |
| 4 | yellow | 835 | SSX2IP |
| 5 | green | 734 | PRB4 |
| 6 | red | 696 | RHOT1 |
| 7 | black | 542 | PCMT1 |
| 8 | pink | 519 | C1orf105 |
| 9 | magenta | 454 | ATP5MC3 |
| 10 | purple | 426 | FUT7 |
| 11 | greenyellow | 395 | HPRT1 |
| 12 | tan | 395 | USP33 |
| 13 | salmon | 390 | RSL24D1 |
| 14 | cyan | 388 | NFAT5 |
| 15 | midnightblue | 358 | TYROBP |

**Figure 3:**
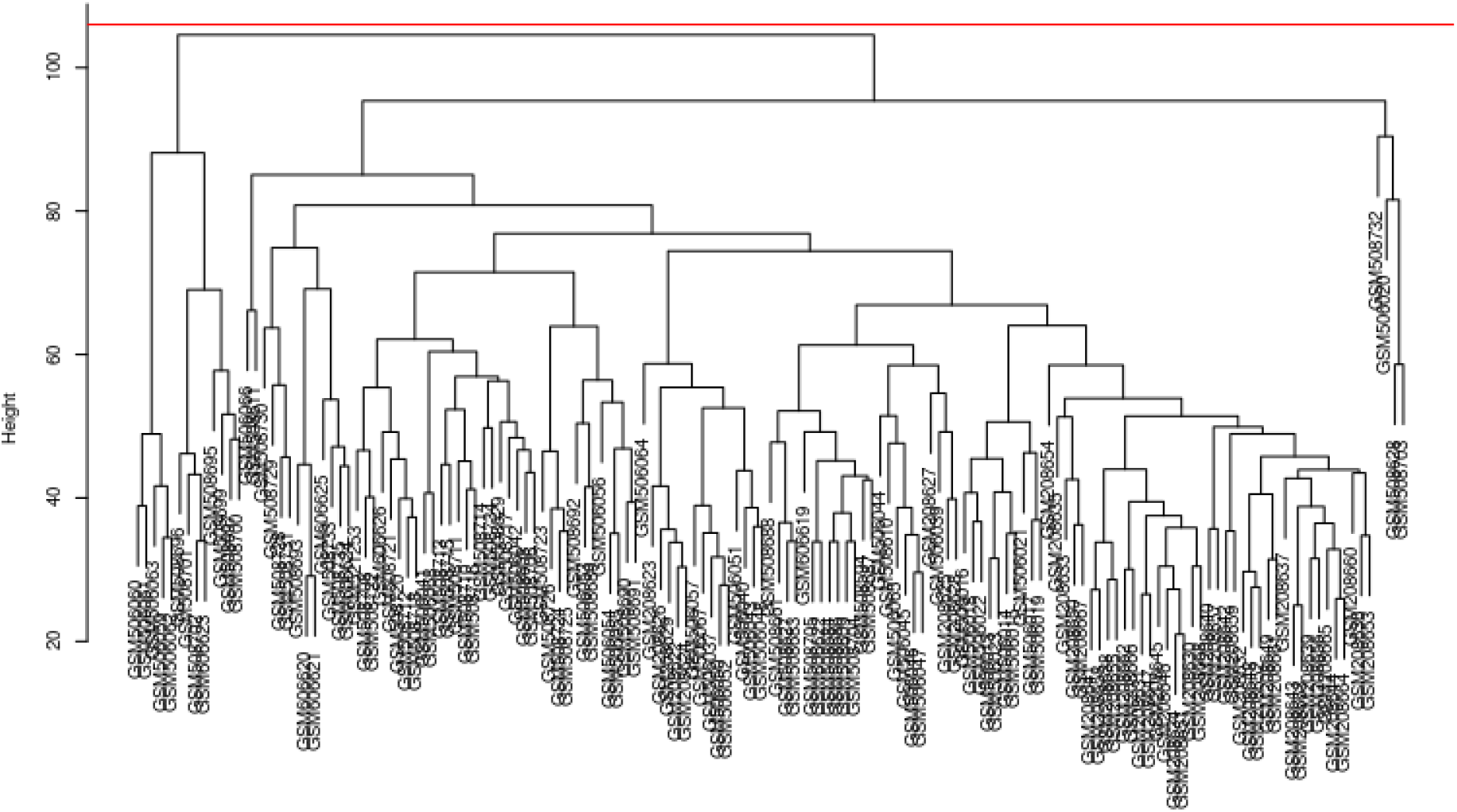
Hierarchical clustering dendrogram based on the sample generated using samples of PD_normal_dataset, the red color line shows an outlier detection threshold at the height of 105. No outlier samples found in PD_normal_dataset.

**Figure 4:**
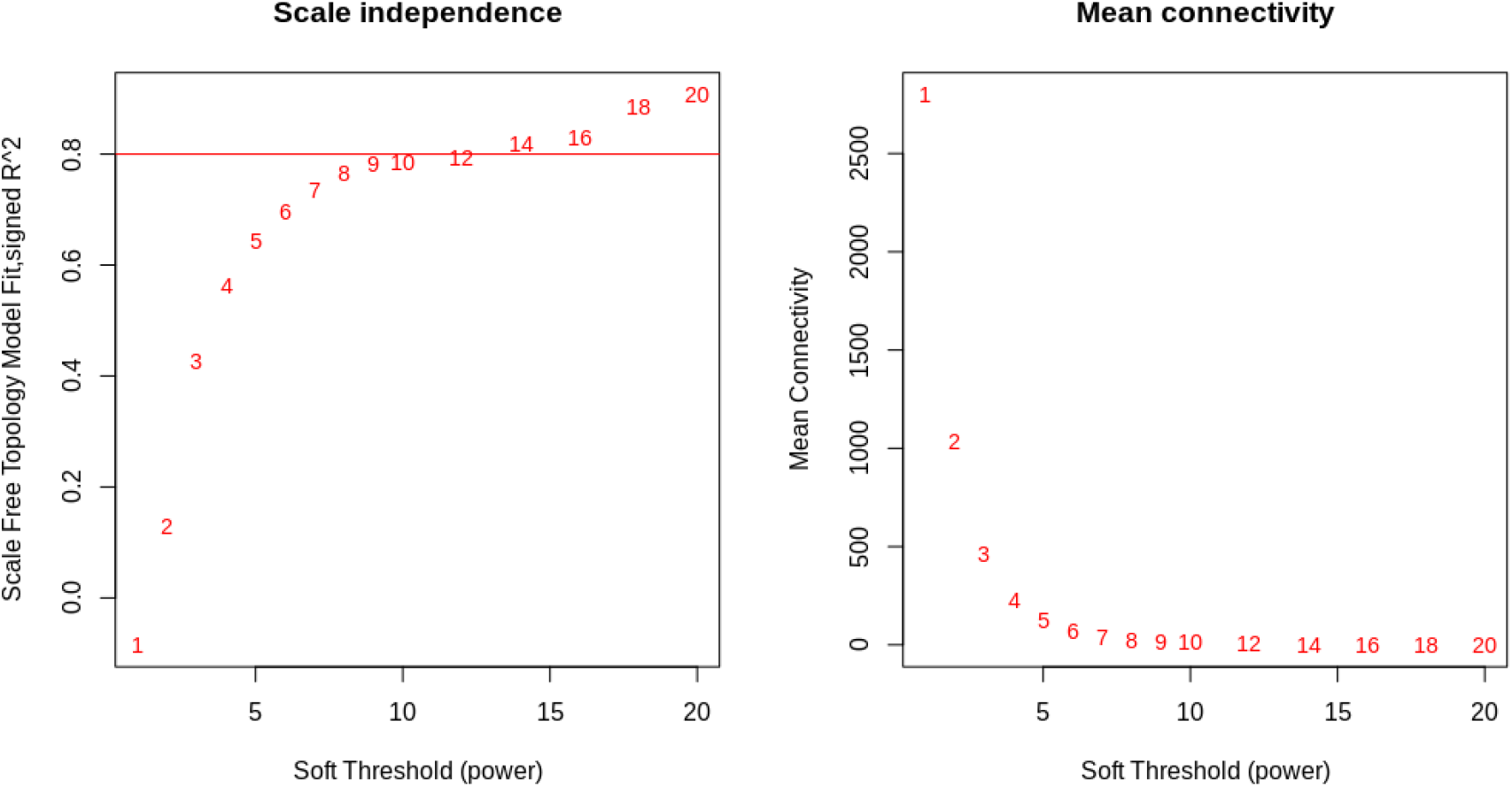
The scale-free index generated using samples of PD_normal_datasetset, the scale-free topology model fit and signed R2 vs. Soft (Power) threshold is shown on the left side, which shows the network achieve scale-free topology at ten. The result on the right side shows the mean connectivity vs. Soft (Power) threshold, which indicates mean connectivity stable at power ten. Both results give ten power, so ten selected as soft thresholds (power).

**Figure 5:**
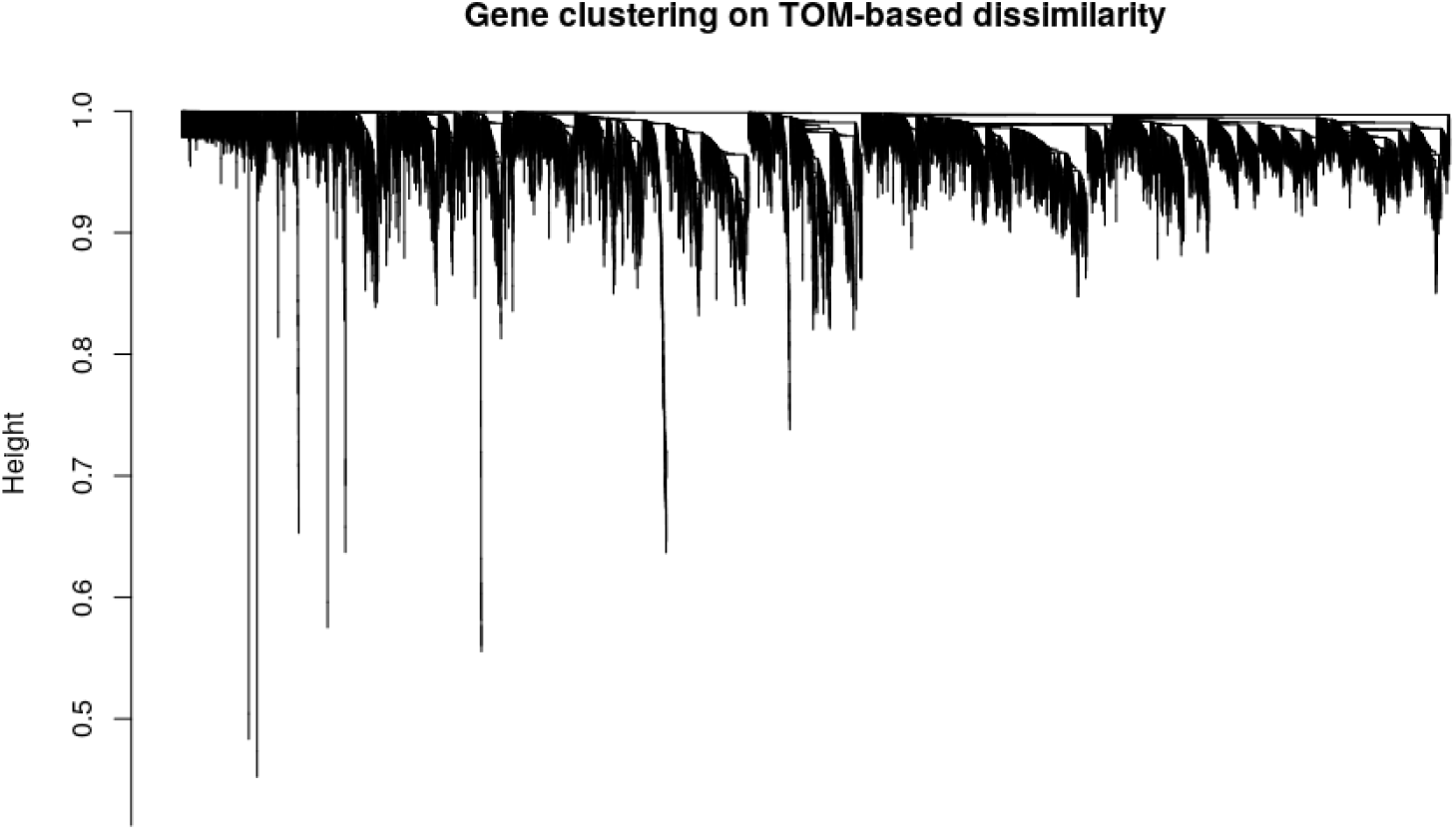
Hierarchical clustering dendrogram obtains using samples of PD_normal_datset, based on the genes using TOM (Yip and Horvath, 2007) based similarity.

**Figure 6:**
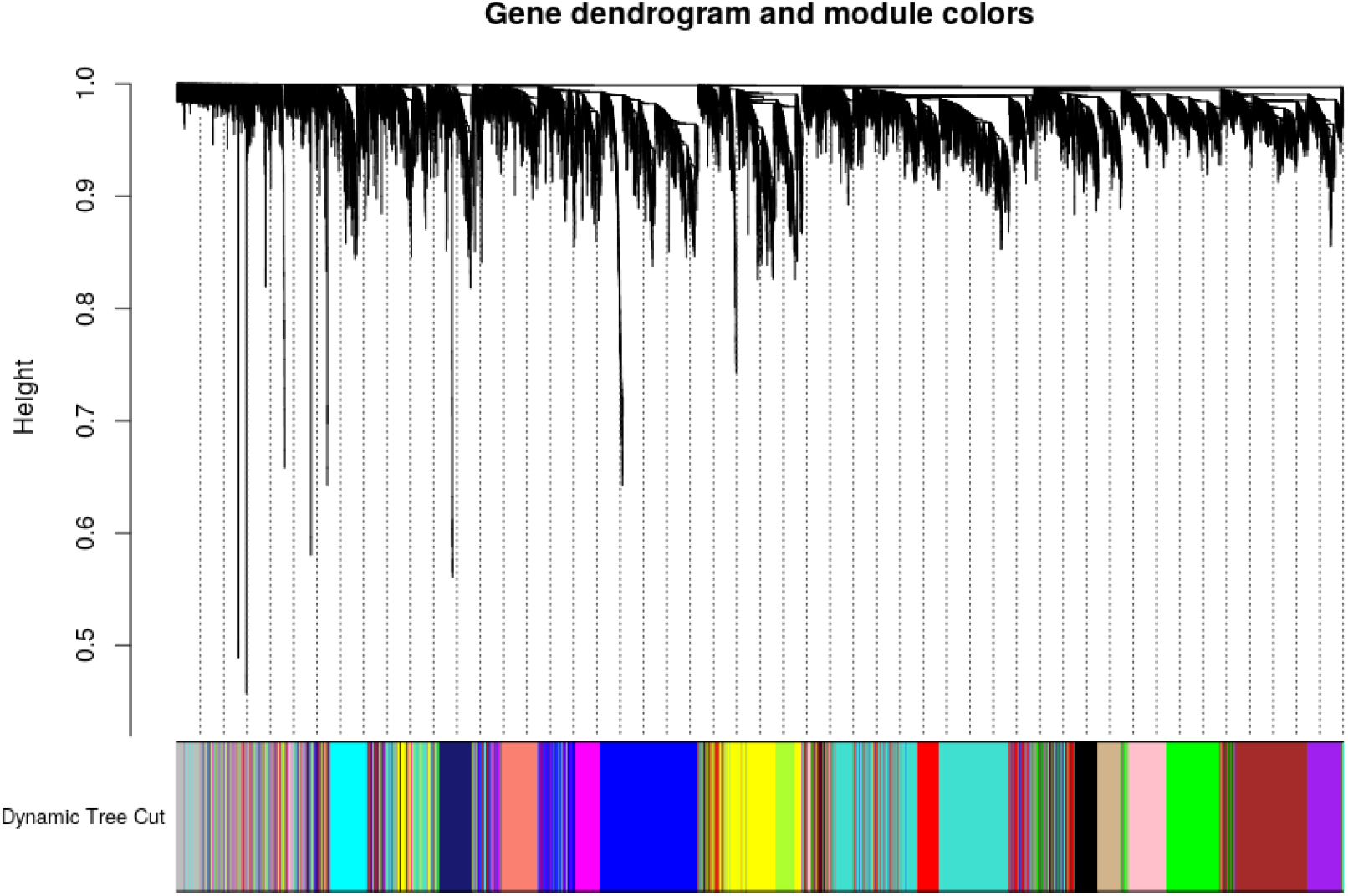
A hierarchical clustering dendrogram generated using samples of PD_normal_dataset based on the genes using TOM (Yip and Horvath, 2007) based similarity with module colors. The individual module represented by colors like turquoise, blue, brown, yellow, green, red, black, and pink. Gray color represents a gene outlier in the dataset.

The correlation between module eigengene (ME) expression value and age, gender (0-male, 1-female), stage (0 - Group1, 1 - Group2 (aging), 2 - PD), PD effect (Group1 and Group2 (aging) - 0, PD - 1) calculated for each module. The module eigengene (ME) expression values of module PD_M4 (Yellow), PD_M6 (Red), PD_M7 (Black), PD_M11 (Greenyellow), and PD_M12 (tan) shows a negative correlation and module PD_M8 (pink), PD_M10 (purple), and PD_M15 (midnight-blue) shows a negative correlation with aging, stage, and PD. The module PD M4 (yellow), PD M6 (red), PD_M7 (black), PD_M9 (magenta), PD_M11 (green-yellow), PD_M12 (tan), and PD_M13 (salmon) show a positive correlation and PD_M3 (brown), PD_M5 (green), PD_M10 (purple) and PD_M14 (cyan) shows a negative correlation with gender. Module PD_11 (green-yellow) and PD 15 (midnight-blue) strongly correlated with both aging and PD and module PD_M6 (red), PD_M7 (black), PD_M10 (purple) and PD_M12 (tan) strongly correlated with only aging. The Figure 7 shows the correlation significance of ME expression values to traits age, gender, stage, and PD_effect.The ME expression values indicate that the genes in PD_M15 module were up-regulated. In contrast, the genes in the PD_M11 module were down-regulated in the transition from Group1 to Group2 (aging) to PD. The Figure 8 shows ME expression values of M11 and M15 module for individual samples according to the grouping Group1, Group2 (aged), and PD.

**Figure 7:**
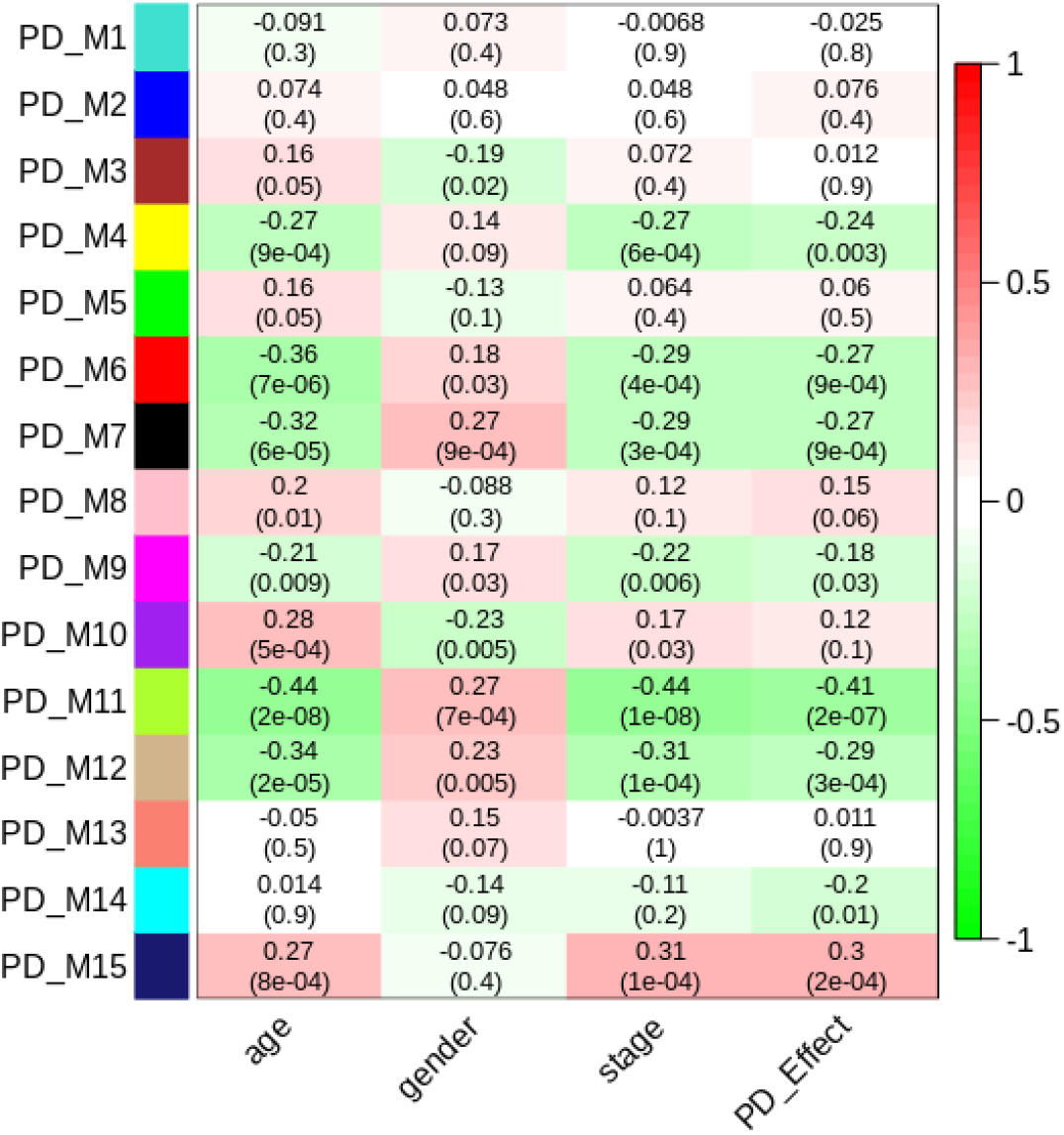
Correlation between ME expression value and value and age, gender (0-male, 1-female), stage (0 - Group1, 1 - Group2 (aging), 2 - PD), PD_effect (Group1 and Group2 (aging) - 0, PD - 1) for each module. Pearson correlation is reported with the p-value given inside the parentheses.

**Figure 8:**
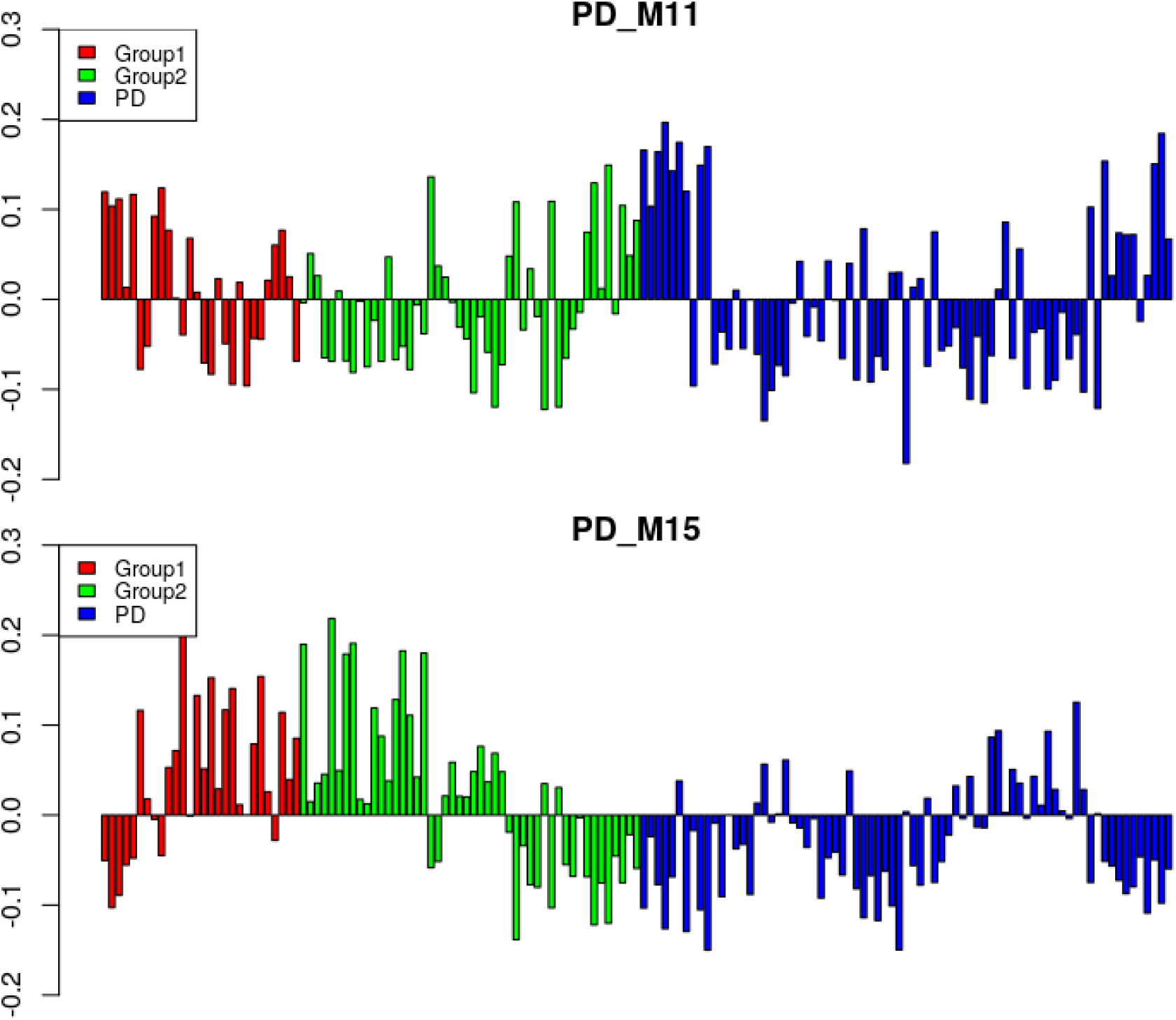
ME expression values of M11 and M15 (y-axis) across samples (x-axis) in PD_normal_dataset. The samples grouped into Group1 (Red), aging (Green), and PD (Blue).

The overlap between the module and cell-type-specific genes was calculated and analyzed. Table 8 represented the p-value obtained from the overlap of the module and cell-type-specific genes. The analysis showed that modules PD_M1 (turquoise), PD_M2 (blue), PD_M4 (yellow), and PD_M13 (salmon) associated with astrocytes cell, modules PD_M4 (yellow), and PD_M11 (green-yellow) associated with neurons cells, module PD_M2 (blue), PD_M11 (green-yellow), PD_M12 (tan), PD_M15 (midnight-blue) associated with endothelial cells, module PD_M6 (red)and PD_M15 (midnight-blue) associated with microglia gene and module PD_M1(turquoise), PD_M4 (yellow) and PD_M13 (salmon) associated with oligodendrocytes cell. On the other hand, modules PD_M3 (brown), PD_M7 (black), PD_M8(pink), PD_M9 (magenta), PD_M10 (purple), and PD_M14 (cyan) showed less significance with the cell type. Further, the module specific to Microglia, neurons, showed a significant correlation with aging or PD.

**Table 8:** Overlap between cell type specific genes and modules

| Module | Astrocytes | Endothelial | Microglia | Neurons | Oligodendrocytes |
| --- | --- | --- | --- | --- | --- |
| PD_M1 | **2.86E-03** | 0.578 | 0.845 | 1.0 | **5.00E-04** |
| PD_M2 | **1.53E-05** | **1.71E-02** | 0.935 | 0.924 | 0.369 |
| PD_M3 | 0.974 | 0.969 | 0.779 | 0.994 | 0.994 |
| PD_M4 | **4.65E-02** | 0.298 | 1.0 | **4.21E-53** | **1.34E-12** |
| PD_M5 | 0.946 | 0.439 | 0.978 | 0.991 | 1.0 |
| PD_M6 | 0.689 | 0.674 | **6.74E-02** | 0.809 | 0.434 |
| PD_M7 | 1.0 | 1.0 | 1.0 | 1.0 | 0.930 |
| PD_M8 | 0.808 | 0.514 | 1.0 | 0.639 | 0.921 |
| PD_M9 | 0.730 | 0.190 | 1.0 | 0.258 | 0.891 |
| PD_M10 | 0.972 | 1.0 | 0.645 | 0.637 | 1.0 |
| PD_M11 | 0.843 | **4.19E-02** | 1.0 | **4.52E-48** | 0.854 |
| PD_M12 | 0.964 | **4.19E-02** | 0.604 | 0.176 | 0.854 |
| PD_M13 | **4.48E-02** | 0.370 | 1.0 | 0.997 | **1.17E-02** |
| PD_M14 | 0.403 | 1.0 | 0.866 | 0.107 | 0.561 |
| PD_M15 | 0.178 | **2.76E-06** | **1.25E-73** | 1.0 | 1.0 |

Module preservation analysis was performed (Langfelder et al., 2011b) using GSE7621, GSE20141, GSE20163, GSE20292 dataset. Most of the aging and PD related modules identified previously showed moderate to high preservation. The modules PD_M1, PD_M3, PD_M7, PD_M9, PD_M11, PD_M12 showed high preservation compared to other modules.The modules specific to neurons (PD_M4 and PD_M11), microglia (PD_M6 and PD_M15), endothelial cells (PD_M11 and PD_M15), and astrocytes (PD_M2, PD_M4,and PD_M13) were preserved in multiple datasets. Since both neuron and glial cells are affected together in PD, we suggest that neuron-glial interactions might be affected in PD. Further, the module PD_M7, PD_M8, PD_M9, and PD_M10, which showed less significance with the cell type, was also preserved with these datasets.The Figure 9,10 showed Z_summary measure of module preservation analysis.

**Figure 9:**
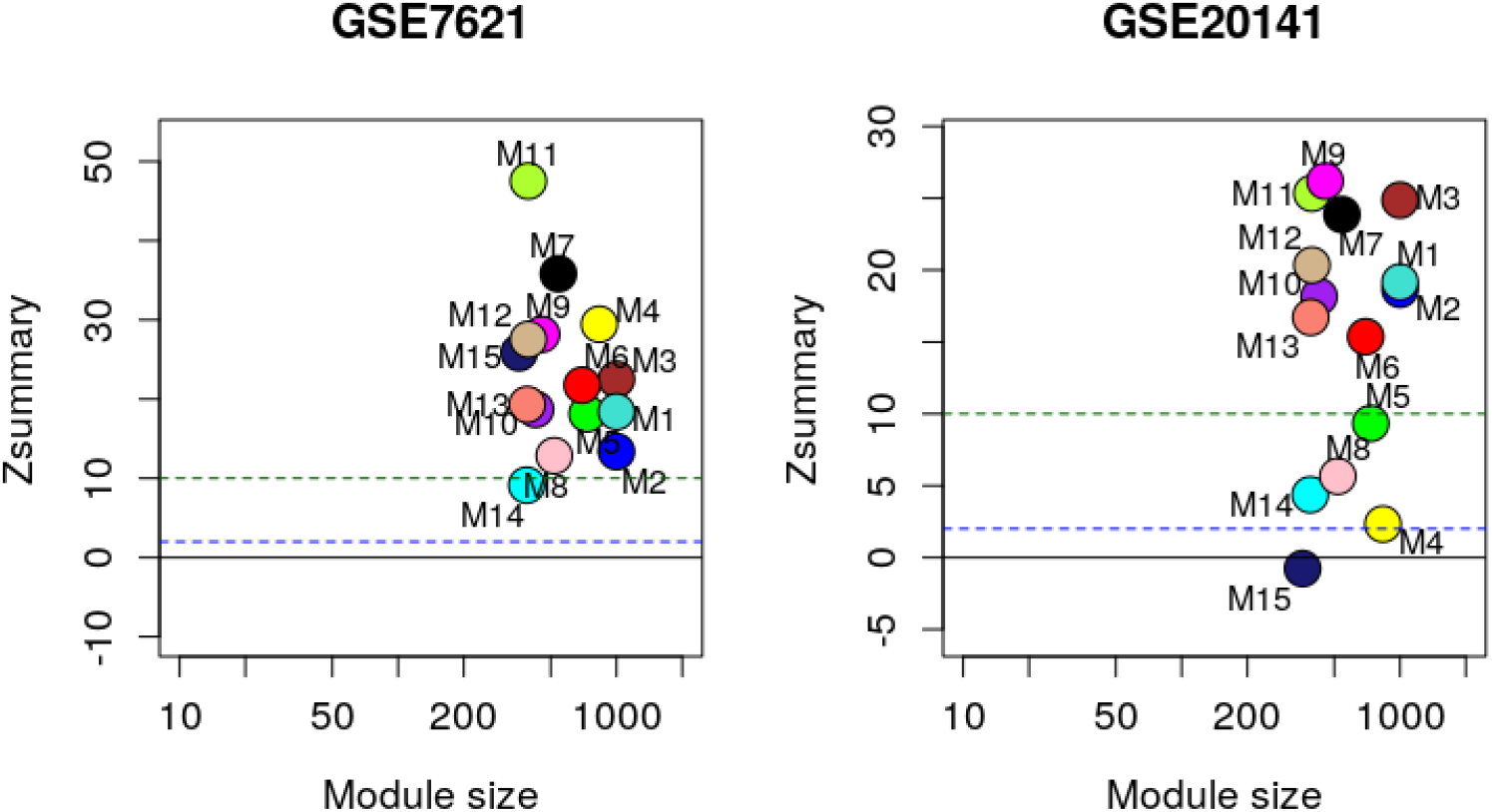
Module preservation analysis using Z_summary with GESE7627 and GSE20141 datasets.

**Figure 10:**
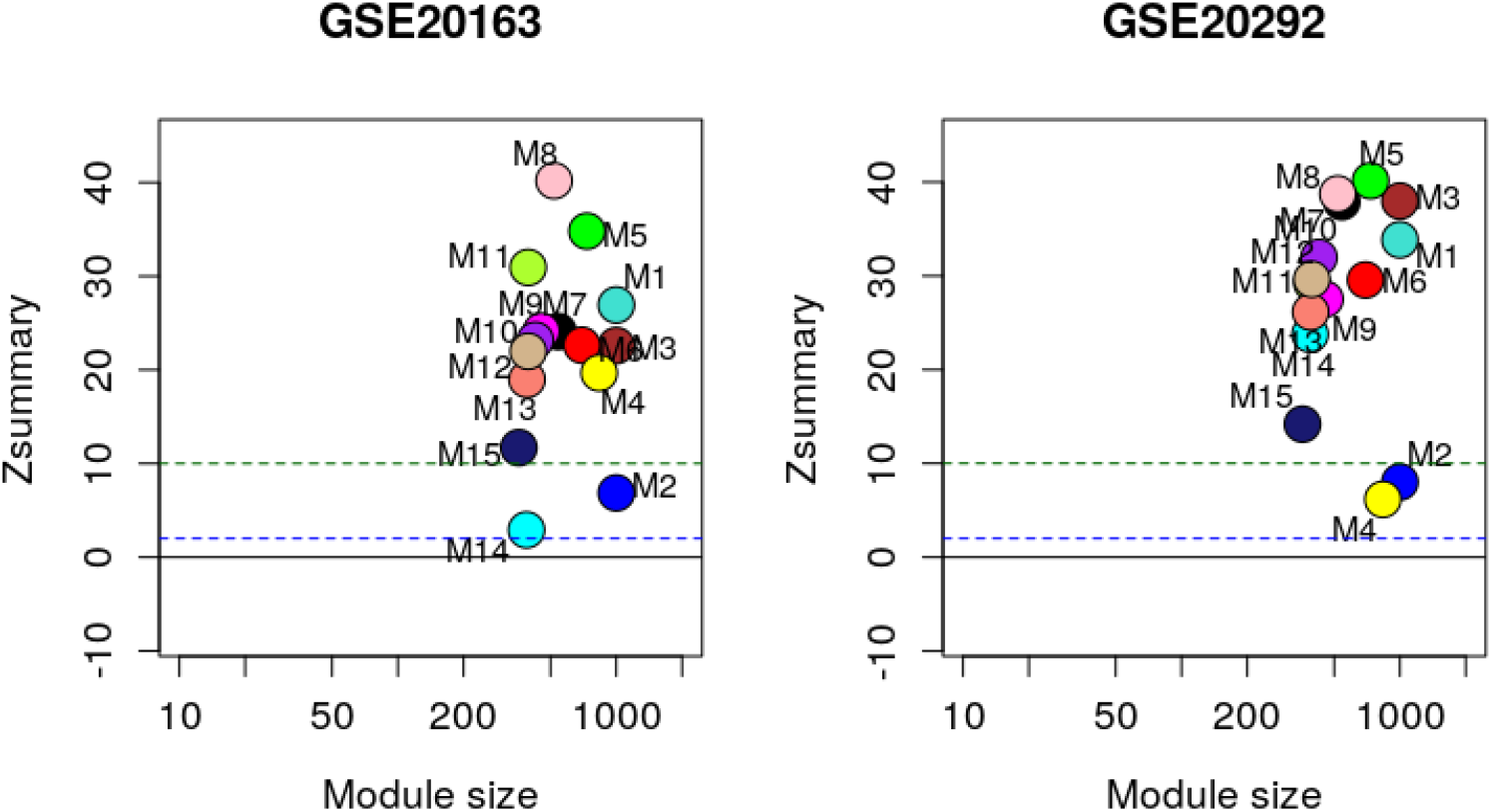
Module preservation analysis using Z_summary with GESE20163 and GSE20292 datasets.

There were several modules that showed significant associations with chronological age and PD and several that showed negative associations were enriched for observing what the genes encoded for.

## 6. DISCUSSION

Using the microarray dataset GSE8397, GSE20164, and GSE20295 from the GeneExpression Omnibus database, which included 75 samples from PD patients and 76 matched controls, 828 DEGs were identified after data preprocessing, WGCNA followed by enrichment analysis and differential expression analysis and its enrichment. The important upregulated and downregulated genes were selected from analysis from all the data (Table 3, 4, 5, 6).

In WCGNA Co-expression analysis obtained by merging the microarray data of three groups viz young, old, and diseased, some biologically meaningful modules were identified. From the heatmap Fig. 7 generated in WGCNA, it was observed that M11 and M15 were two modules that showed the highest p-value. Of this M11 had the highest negative correlation and M15 had the highest positive correlation in context to aging and PD effect. The enrichment analysis study done on M11 and M15 clusters (Table 9, 10) showed the following result.

**Table 9:**
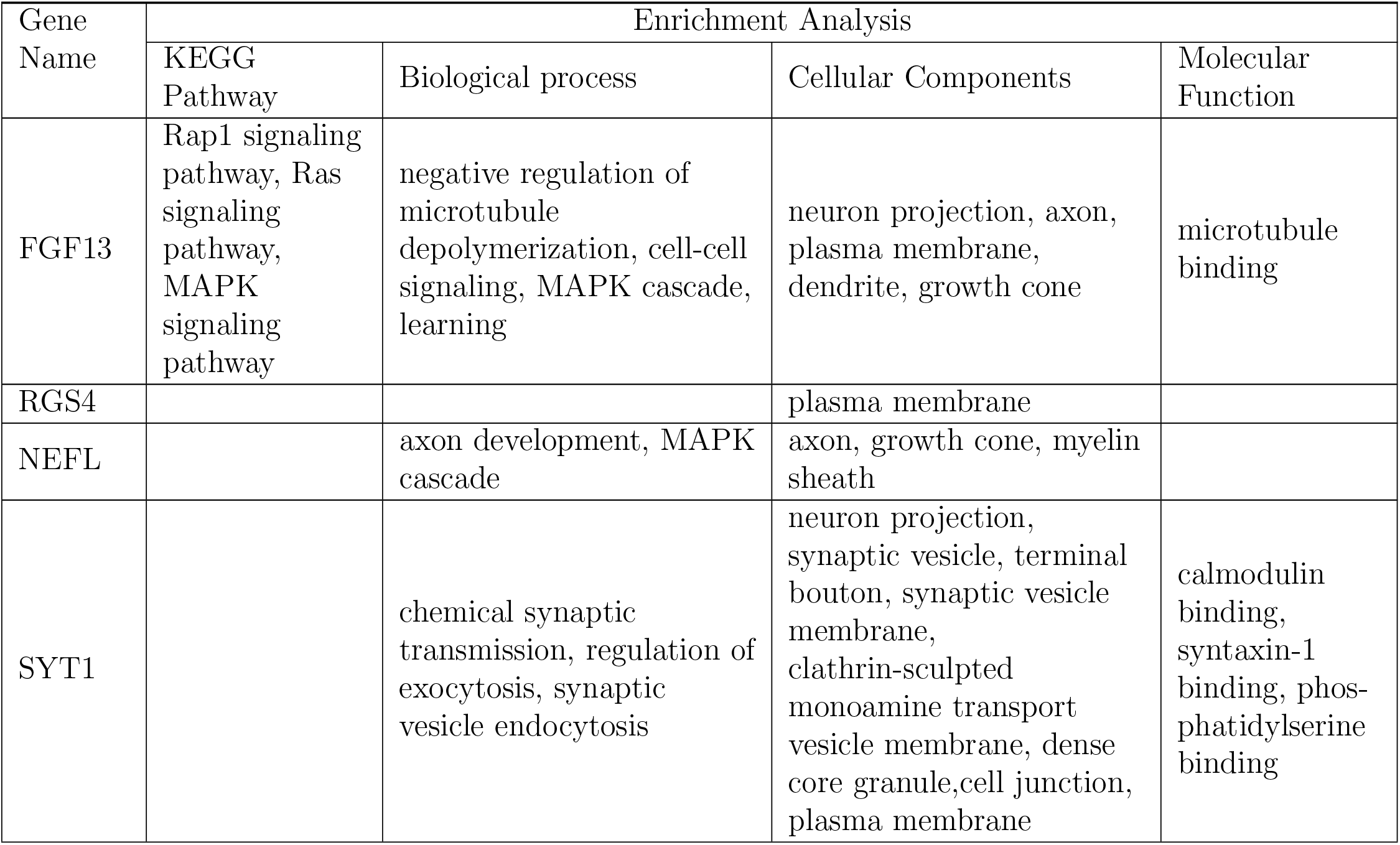
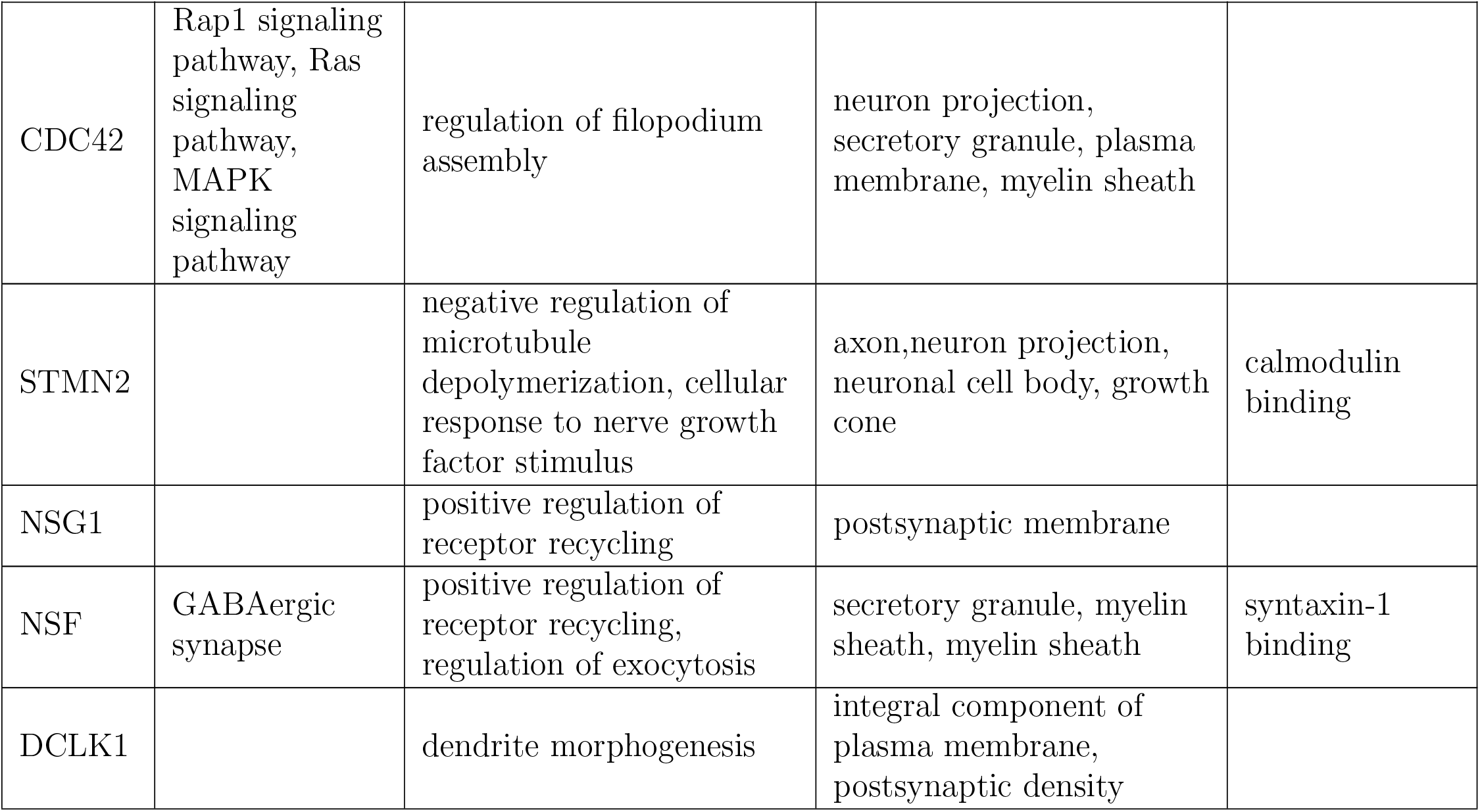
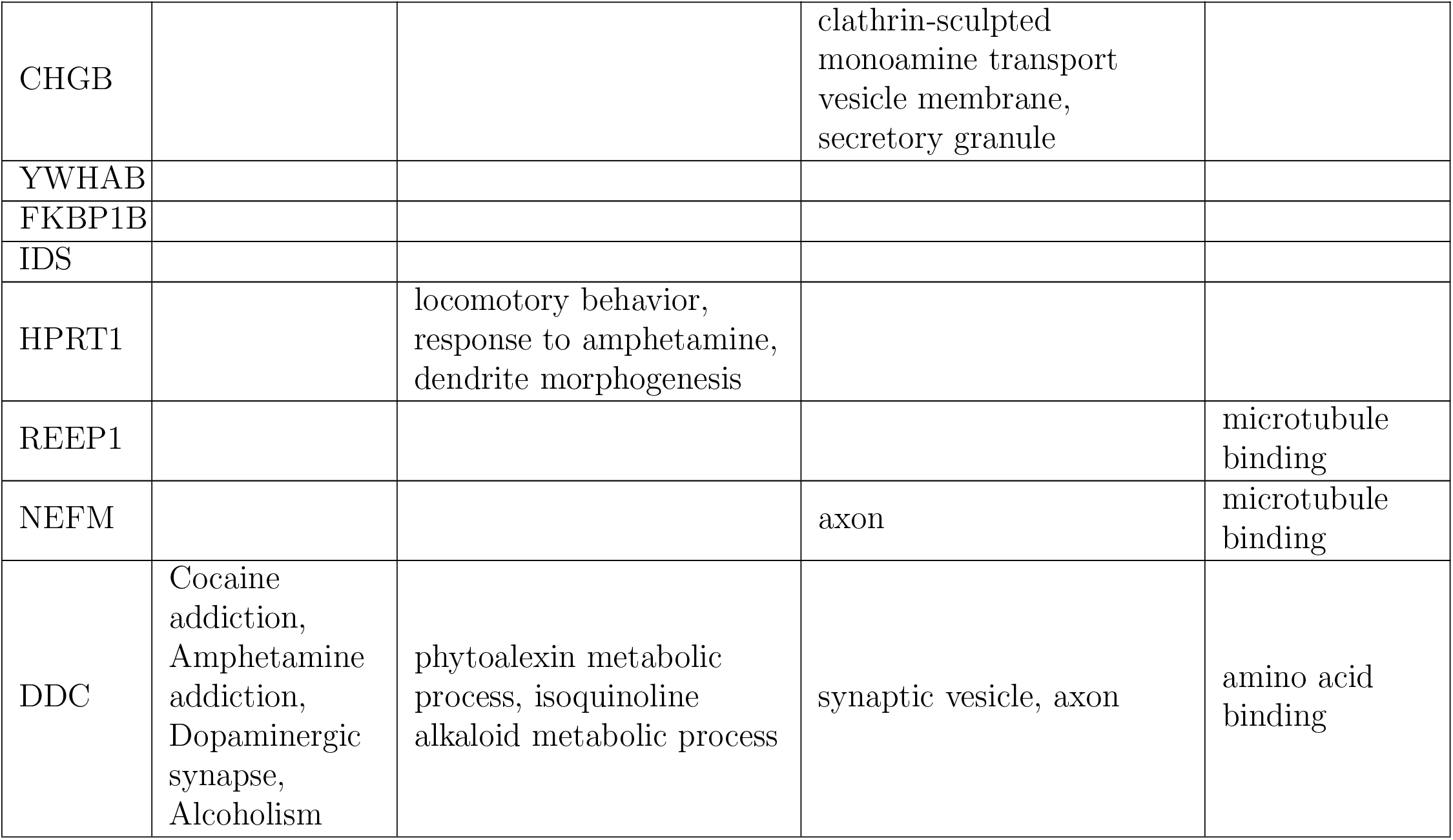
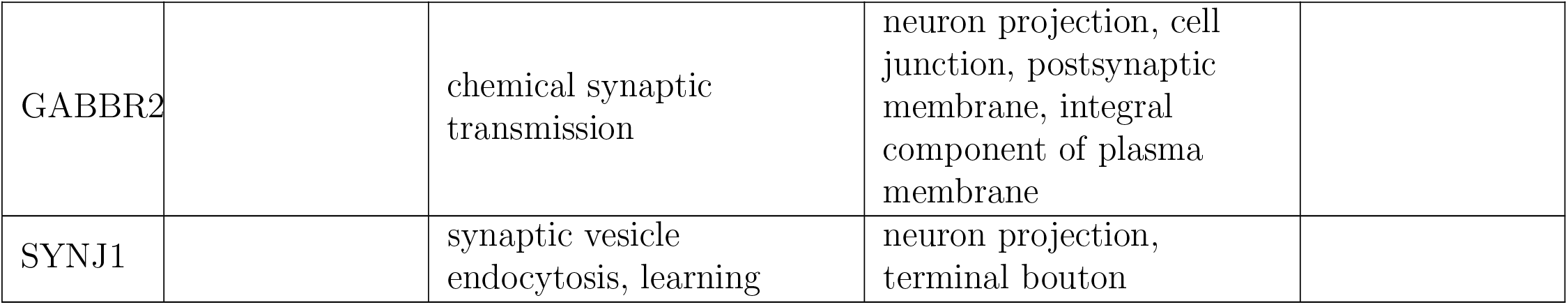
Detail Enrichment analysis of 19 downregulated genes common in Cluster 11 and DEG analysis.

**Table 10:**
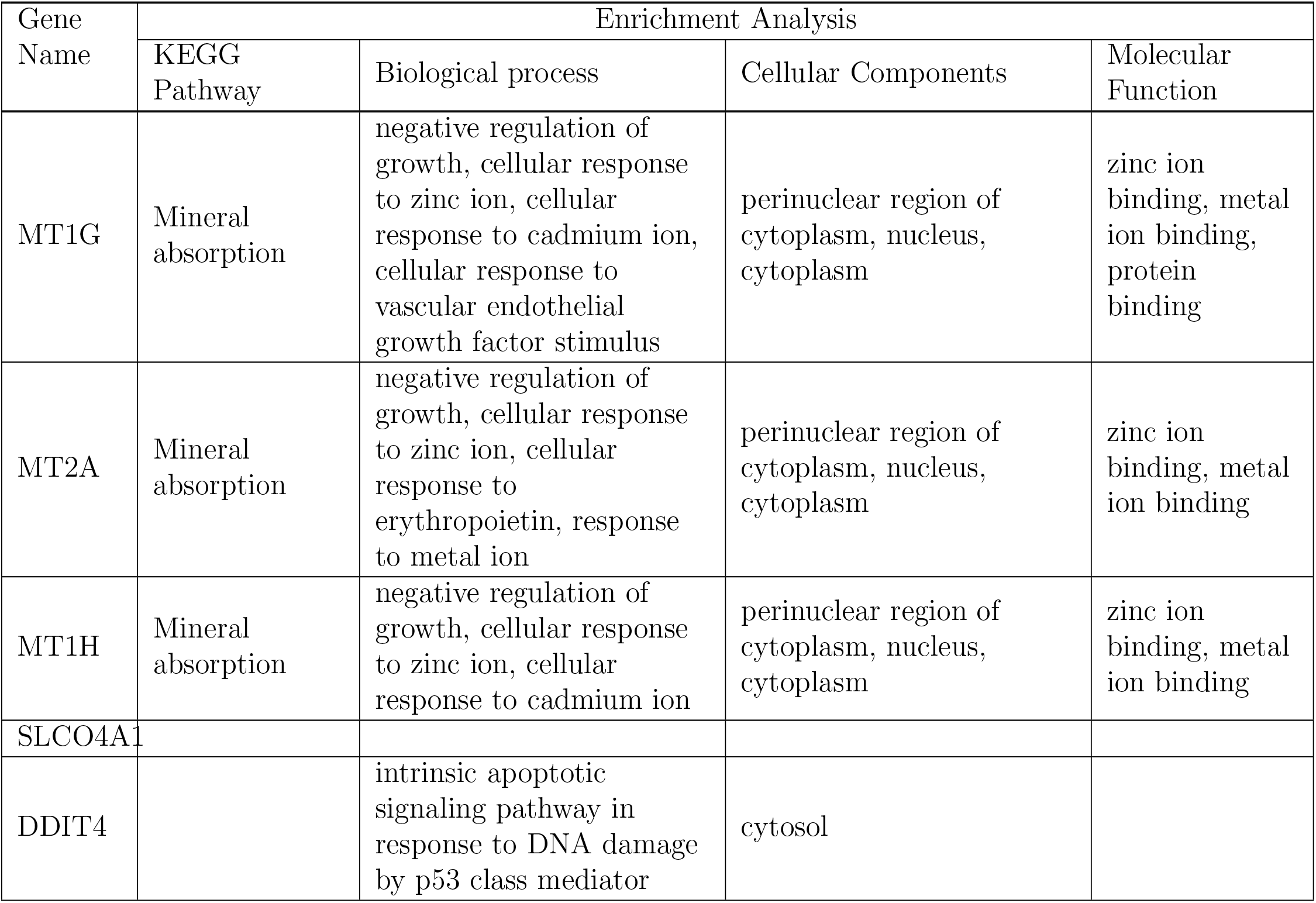
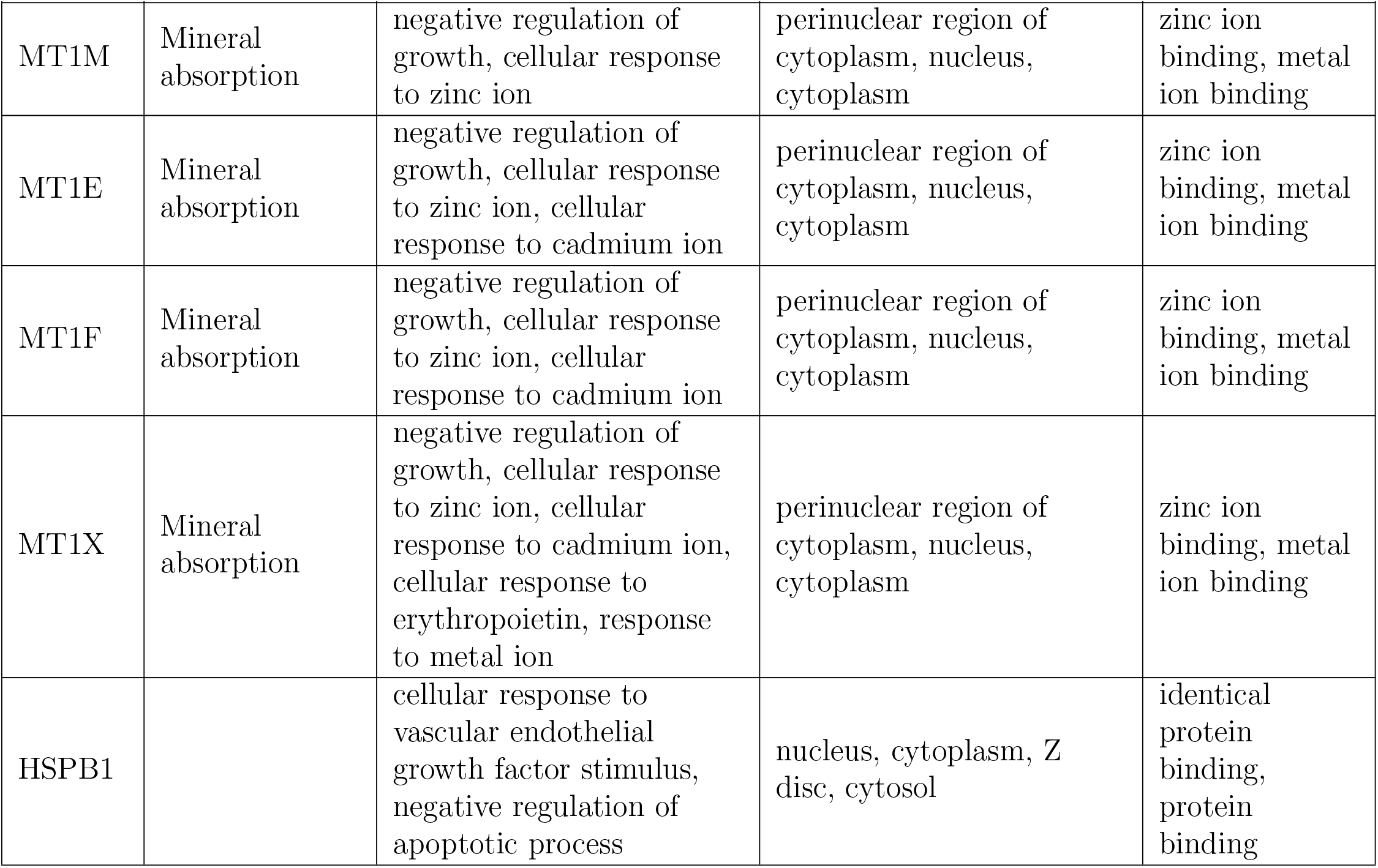
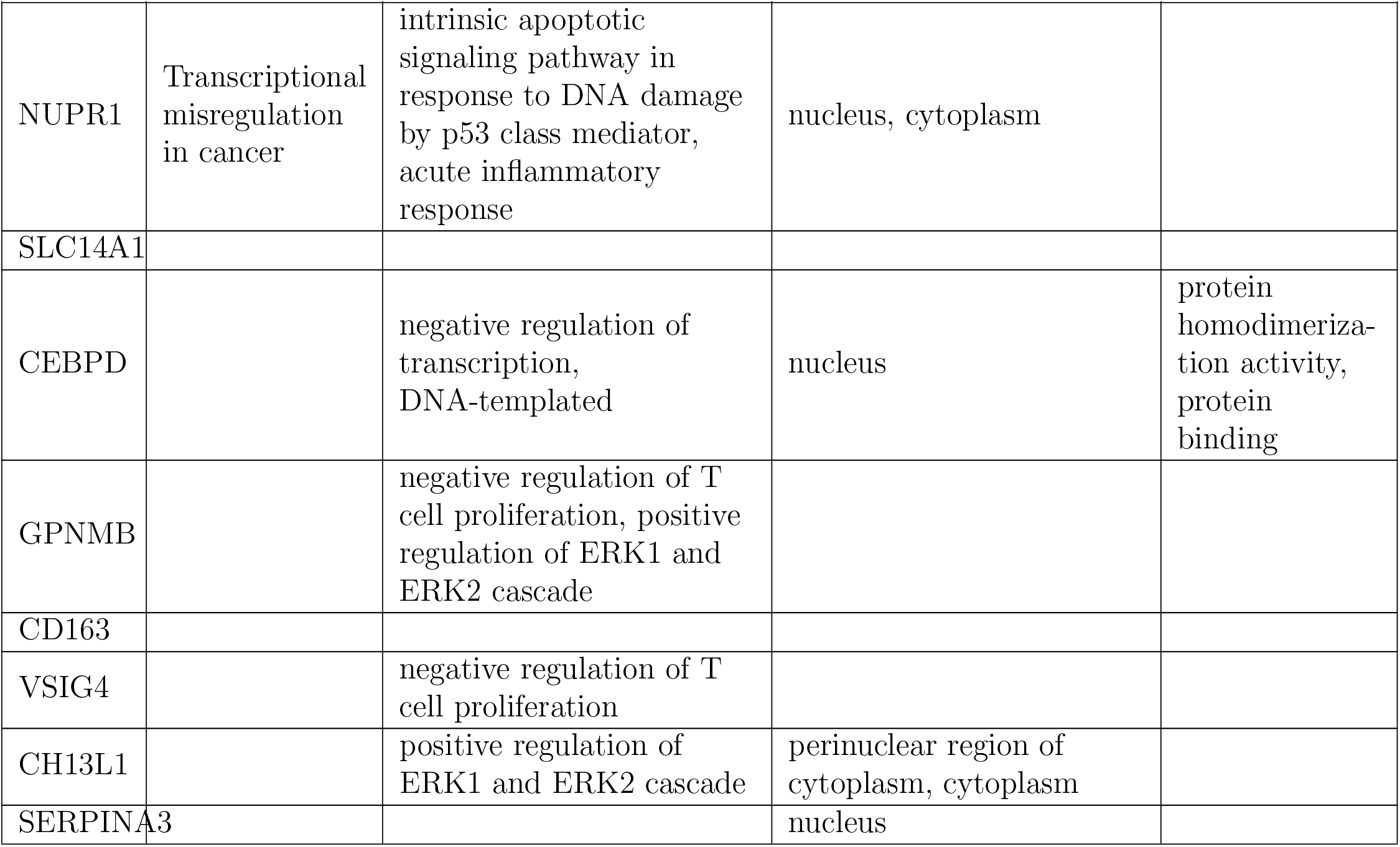
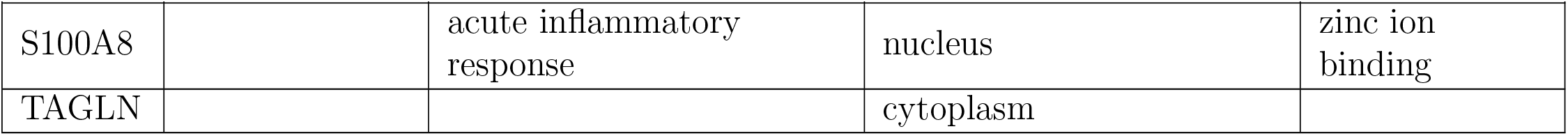
Detail Enrichment analysis of 20 upregulated genes.

| Gene Name | Enrichment Analysis |  |  |  |
| --- | --- | --- | --- | --- |
|  | KEGG Pathway | Biological process | Cellular Components | Molecular Function |
| MT1G | Mineral absorption | negative regulation of growth, cellular response to zinc ion, cellular response to cadmium ion, cellular response to vascular endothelial growth factor stimulus | perinuclear region of cytoplasm, nucleus, cytoplasm | zinc ion binding, metal ion binding, protein binding |
| MT2A | Mineral absorption | negative regulation of growth, cellular response to zinc ion, cellular response to erythropoietin, response to metal ion | perinuclear region of cytoplasm, nucleus, cytoplasm | zinc ion binding, metal ion binding |
| MT1H | Mineral absorption | negative regulation of growth, cellular response to zinc ion, cellular response to cadmium ion | perinuclear region of cytoplasm, nucleus, cytoplasm | zinc ion binding, metal ion binding |
| SLCO4A1 |  |  |  |  |
| DDIT4 |  | intrinsic apoptotic signaling pathway in response to DNA damage by p53 class mediator | cytosol |  |

Continue on the next page
| Gene Name | Enrichment Analysis |  |  |  |
| --- | --- | --- | --- | --- |
|  | KEGG Pathway | Biological process | Cellular Components | Molecular Function |
| MT1M | Mineral absorption | negative regulation of growth, cellular response to zinc ion | perinuclear region of cytoplasm, nucleus, cytoplasm | zinc ion binding, metal ion binding |
| MT1E | Mineral absorption | negative regulation of growth, cellular response to zinc ion, cellular response to cadmium ion | perinuclear region of cytoplasm, nucleus, cytoplasm | zinc ion binding, metal ion binding |
| MT1F | Mineral absorption | negative regulation of growth, cellular response to zinc ion, cellular response to cadmium ion | perinuclear region of cytoplasm, nucleus, cytoplasm | zinc ion binding, metal ion binding |
| MT1X | Mineral absorption | negative regulation of growth, cellular response to zinc ion, cellular response to cadmium ion, cellular response to erythropoietin, response to metal ion | perinuclear region of cytoplasm, nucleus, cytoplasm | zinc ion binding, metal ion binding |
| HSPB1 |  | cellular response to vascular endothelial growth factor stimulus, negative regulation of apoptotic process | nucleus, cytoplasm, Z disc, cytosol | identical protein binding, protein binding |

Continue on the next page
| Gene Name | Enrichment Analysis |  |  |  |
| --- | --- | --- | --- | --- |
|  | KEGG Pathway | Biological process | Cellular Components | Molecular Function |
| NUPR1 | Transcriptional misregulation in cancer | intrinsic apoptotic signaling pathway in response to DNA damage by p53 class mediator, acute inflammatory response | nucleus, cytoplasm |  |
| SLC14A1 |  |  |  |  |
| CEBPD |  | negative regulation of transcription, DNA-templated | nucleus | protein homodimerization activity, protein binding |
| GPNMB |  | negative regulation of T cell proliferation, positive regulation of ERK1 and ERK2 cascade |  |  |
| CD163 |  |  |  |  |
| VSIG4 |  | negative regulation of T cell proliferation |  |  |
| CH13L1 |  | positive regulation of ERK1 and ERK2 cascade | perinuclear region of cytoplasm, cytoplasm |  |
| SERPINA3 |  |  | nucleus |  |
|  | KEGG Pathway | Biological process | Cellular Components | Molecular Function |
| S100A8 |  | acute inflammatory response | nucleus | zinc ion binding |
| TAGLN |  |  | cytoplasm |  |

- The top 5 significantly enriched GO terms in biological processes are axon development, protein localization to plasma membrane, neurofilament bundle assembly, exocytsis, and inositol phosphate metabolic process for DEG IN M11.
- The top5 significantly enriched molecular function annotations for genes differentially ex-pressed in M11 are protein binding, structural constituent of cytoskeleton, SNARE binding, protein kinase binding, and ATP binding.
- The top 5 significantly enriched cellular component annotations for genes differentially expressed in M11 are myelin sheath, cytosol, synaptic vesicle, and synaptic vesicle membrane, mitochondrion and neuron projection and neurofilament.
- The top 5 significantly enriched GO terms in biological processes for M15 are : The top 5 significantly enriched GO terms in biological processes are inflammatory response, immune response,innate immune response, interferon gamma mediated signalling pathway, cellular response to lipopolysaccharide. The Top 5 significantly enriched molecular function annotation for DEG in M15 are MHC class II receptor activity,protein binding, peptide antigen binding,IgG binding, receptor activity. The top 5 significantly enriched cellular component annotations for DEG in M15 are extracellular exosome, MHC class II protein complex, plasma membrane, extracellular space, integral component of plasma membrane.

An overlap between cell type specific genes and modules (Table 8) showed that Module M11 is associated with neurons and endothelial cells, M15 had wide association with microglia.

From the differential Expression Analysis, using volcano plot (Fig. 2), 568 upregulated and 126 downregulated genes were identified, from which the top 25 genes (from each cluster) were ranked using logFC.

The data obtained in DEG analysis had three clusters viz Group 1 vs PD, Group 1 vs Group 2 and Group 2 vs PD, taking into account the parameters of influence of ageing on PD (Table 3, 4, 5, 6).

On close study of comparison between Group1 vs PD along with Group 1 vs Group 2, an assessment of the differential expression in context to both aging and PD was done. It has been noticed that genes that showed downregulation/upregulation for both PD samples and Group 2 are more likely to reflect age dependant contribution to the disease risk rather than genes altered only in PD (Wood-Kaczmar et al., 2006). Parkinson’s Disease is known to be an age related neurodegenerative disease (Nussbaum and Ellis, 2003). It affects 41 people per 100,000 in the age group of 30-40 years old to over 1900 per 100,000 in people over 80 years of age, thus showing its high prevalence in senior population (Lau and Breteler, 2006). Table refEnrichment analysis 19 downregulated genes, 10 summarizes those genes that are common in both the clusters i.e Group 1 vs PD and Group1 vs Group 2.

A combined analysis of the enrichment data (using DAVID 6.7) for both upregulated and down-regulated genes obtained by merging the two different methods (WGCNA and differential expression analysis) narrowed down the number of significant DEG to 39. It was observed that the list of DEG downregulated (Group1 vs Group2, Group 1 vs PD and Group2 vs PD) from DEG analysis had 19 genes common with the Cluster 11 module of WGCNA. These genes were FGF13, RGS4, NEFL, SYT1, CDC42, STMN2, NSG1, NSF, DCLK1, CHGB,YWHAB, FKBP1B, IDS, HPRT1, REEP1, NEFM, DDC, GABBR2, and SYNJ1. Similarly, a list of DEG (Group 1 vs PD and Group2 vs PD) from DEG analysis had 20 upregulated genes in common with the M15 cluster of WGCNA study. These were MT1G, MT2A, MT1H, SLCO4A1, DDIT4,MT1M, MT1E, MT1F, MT1X, HSPB1, NUPR1, SLC14A1, CEBPD, GPNMB, CD163,VSIG4, CH13L1, SERPINA3, S100A8, TAGLN.Table 9, 10 shows details of Enrichment analysis of 19 downregulated genes and 20 downregulated genes.

## 7. Conclusion

Parkinson’s Disease has been considered as a disorder of cell metabolism (Anandhan et al., 2017). Some of the metabolic pathways that result in the neurological disorder have been identified. Study of the correlations between PD using gene expression comparisons between normal and diseased human samples can lead to identifying important genes and pathways which can contribute to the process of disease diagonosis, study of disease progression, identification of biomarkers and drug discovery. Among the various pathways, alterations in redox homeostasis and bioenergetics are thought to be the central component of neurodegeneration that contribute to the impairment of important homeostatic process in dopaminergic cells (Anandhan et al., 2017). Similarly, GABA plays an important role in behaviour, cognition, and the body’s response to stress (Jie et al., 2018). GABA is the major inhibitory neurotransmitter in the Central Nervous System. The long term stability and function of the neuronal network is dependent on a maintained balance between excitatory and inhibitory synaptic transmission (Tatti et al., 2017). The purpose of our work was to identify such genes that are essentially involved in bringing alterations at various levels of the disease progression. For this, our study involved analysis of gene expression data (microarray data) to cluster the genes on the basis of their expression as suggested from previous works by (Papapetropoulos et al., 2006). The important genes were identified that are significantly upregulated or downregulated with ageing and/or onset of the disease.The function of these potential genes was then checked through the annotations available by enrichment analysis from available gene annotation databases like KEGG (Kanehisa, 2000). Further correlation to the disease could then be interpreted.

From comparative study of the WGCNA and DEG analysis results, it was observed that the top 25 downregulated and upregulated genes obtained in DEG analysis (Table3, 4, 5, 6) had maximum overlap with the two modules viz. Cluster 11 and Cluster 15 of WGCNA clusters obtained. These two clusters had shown highest negative and positive correlation respectively with respect to ageing and PD as seen in the heatmap shown in Fig. 7. This overlapping of a major number of genes, showed the following conclusions:-

- WGCNA method gave robust clusters in this study.
- The overlapping or common genes from the two studies could be marked as potential risk genes in relation to PD, on the basis of their

- statistical significance of upregulation or down regulation of expression against healthy sample.
- Enrichment analysis study of these genes.
- literature surveys on reported correlation of these genes with PD.

Some of the important genes that were identified as important downregulated genes in relation to both ageing and PD, as studied in all the methods viz. WGCNA and DEG analysis are SYT1, NEFL, RGS4, CDC42, FGF13, CHGB, NSG1, DCLK1. These genes have been previously reported to be associated with PD (Sonntag et al., 2009; Su et al., 2018; Sakharkar et al., 2019b; Magalingam et al., 2015). Similarly, among the upregulated genes, VSIG4, MT1H, SLCO4A1, CEBPD, CD163, GPNMB,(Sakharkar et al., 2019b; Li et al., 2019) were previously reported to be associated with PD.

A cell type overlap study with the two important modules M11 and M15 shows the highest association with neuron and microglial cells respectively(Table 8).This shows that most of the downregulated gene expressions have occurred in the neuron cells while the maximum upregulation of gene expression has been found in the microglial cells. This can be assessed as the presence of significantly downregulated genes in the neuron cells and upregulated genes in the microglial cells. Prominent down-regulation of members of the PARK gene family and dysregulation of multiple genes associated with programmed cell death and survival, deregulation of genes for neurotransmitter and ion channel receptors have been reported in several studies including that of Simunovic etal (Simunovic et al., 2008). The present study reported here shows similar association of gene expression changes in particular cell types.

Thus, this study identified a set of potential crucial PD associated genes which were associated with neuron development and differentiation.By implementing both the methods viz WGCNA and DEG analysis, putative risk genes could be identified from the microarray data of the brain region on the basis of gene upregulation or downregulation with respect to young and healthy samples. Further, the comparative analysis from enrichment analysis study of both the methodologies indicate that these genes are involved in particular biological processes, cellular components, and molecular functions that contribute to the motor and non-motor symptoms of the disease. Of the 568 upregulated and 126 downregulated genes, 19 downregulated genes were found to play significant roles in various signalling pathways (FGF13, CDC42), axon development (NEFL), chemical synaptic transmission(SYT1), neuron projection(FGF13), calmodulin binding (RGS4),regulation of exocytosis (SYT1). Similarly, 20 upregulated genes were found to be enriched in several processes like metal binding proteins associated with neuroprotecytion (MT1G, MT2A, MT1H, MT1M, MT1E, MT1F, MT1X),stress responsive genes related to regulation of cellular reactive oxygen species(DDIT4),transporter family (SLCO4A1, SLC14A1),transcriptional regulator (NUPR1, CEBPD),negative regulation of T cell proliferation (VSIG4), inflammatory response (TAGLN,CD163, CH13L1). It was also observed from the WGCNA modules heatmap generated using several parameters (Fig. 7), that the differential gene expression related to PD and ageing were related, confirming age to be a major risk factor for PD (Dawson and Dawson, 2003). Previous studies have shown that the neuron cells show high amount of downregulated gene expressions while the microglial cells show higher upregulation in gene expressions in PD (Sonninen et al., 2020). The identified potential risk genes could be further compared to microarray data from blood samples of healthy and PD individuals to suggest tentative biomarkers in disease diagonosis, disease progression, effect of treatment and therapy.

## Footnotes

1 https://www.ncbi.nlm.nih.gov/geo/

## Notes

### Competing Interest Statement

The authors have declared no competing interest.

